# Acute EIF4G1 depletion reveals transcript-wide determinants of initiation factor dependence and activates initiation-specific ribosome quality control

**DOI:** 10.64898/2026.04.17.719311

**Authors:** Zachary D. Knotts, Michael T. Marr

**Author notes:** Corresponding author: Michael T. Marr II.

## Abstract

Canonical cap-dependent translation initiation requires the eIF4F complex, in which the scaffold protein EIF4G1 bridges cap-bound EIF4E, the RNA helicase EIF4A, and ribosome-associated EIF3. Using an inducible degron to acutely deplete EIF4G1 in human cells, we combined polysome profiling with mRNA-seq to quantify transcript-specific relative polysome association (RPA). EIF4G1 depletion causes heavy polysome collapse with reciprocal 80S monosome accumulation and activates initiation-specific ribosome quality control (iRQC), evidenced by RPS3 mono-ubiquitination enriched on 80S monosomes. Using RPA analysis we find the ribosome association of most transcripts is unaffected. However, we identify 1,354 mRNAs that are Sensitive to EIF4G1 loss and 1,844 mRNAs that are translationally Resistant under the same conditions. As a class, the Resistant transcripts tend to be longer and have a higher GC content in the coding sequence relative to Sensitive or unaffected transcripts. A synthetic reporter confirms that elevated CDS GC content can promote translation upon EIF4G1 depletion. These results suggest that the architecture of the entire transcript, beyond the 5′UTR, affects EIF4G1 usage and that removing EIF4G1 triggers iRQC consistent with an altered initiation architecture.

## Introduction

Canonical cap-dependent translation in eukaryotes begins with the assembly of the eIF4F cap-binding complex at the 7-methylguanosine (m⁷G) cap. Subsequently, the recruitment of the 43S pre-initiation complex, ribosome scanning, and start-codon selection allow protein synthesis to initiate (reviewed in (Brito Querido et al. 2024). Within eIF4F, EIF4G1 serves as a central scaffold bringing together multiple players in translation initiation. EIF4G1 is a large multidomain protein with binding sites for multiple factors involved in translation initiation as well as RNA. It bridges cap-bound EIF4E and the helicase EIF4A to 40S-bound EIF3 and engages poly(A)-binding protein (PABP) bound at the 3′ poly(A) tail (Brito Querido et al. 2024). Through these interactions, EIF4G1 is thought to promote functional mRNA circularization, stabilize initiation complexes, enhance ribosome loading (Hinnebusch 2014), and allow for efficient ribosome recycling (Pisarev et al. 2007). Loss or inhibition of eIF4F components typically depresses global protein synthesis, yet cells often retain translation of specific transcripts, implying factor-specific rules of dependence (Larsson et al. 2007; Hsieh et al. 2012; Thoreen et al. 2012; Truitt et al. 2015; Haizel et al. 2020).

Classically, eIF4F dependence has been attributed to properties of the 5′ untranslated region (5′UTR). Highly structured 5′UTRs, upstream AUGs, and regulatory motifs increase the need for EIF4A1/EIF4G1-driven remodeling (Pause et al. 1994; Svitkin et al. 2001; Iwasaki et al. 2016; Waldron et al. 2019), whereas IRES-like elements or specialized leaders can reduce it (Logan and Shenk 1984; Jang et al. 1988; Pelletier and Sonenberg 1988). mTOR signaling further modulates eIF4F complex formation, creating transcript-selective effects, for example on mRNAs containing a terminal oligopyrimidine (TOP) motif (Thoreen et al. 2012). Although this 5′UTR-centric view explains many observations, it does not fully account for the breadth of responses seen when the eIF4F scaffold is perturbed (Thoreen et al. 2012; Lee et al. 2016; Lahr et al. 2017).

The eIF4G family in mammals includes two additional members. EIF4G2, also known as DAP5 or NAT1, lacks the N-terminal domains responsible for binding EIF4E and PABP (Imataka et al. 1998). Despite this, DAP5 promotes translation of specific mRNA subsets, particularly those with structured 5′UTRs or upstream open reading frames, through cap-independent or eIF3d-mediated mechanisms (de la Parra et al. 2018; Haizel et al. 2020; Weber et al. 2022).

Multiple studies have collectively identified several hundred DAP5-dependent transcripts across diverse cell types and conditions (Liberman et al. 2009; Yoffe et al. 2016; de la Parra et al. 2018; Volta et al. 2021; David et al. 2022; Weber et al. 2022). EIF4G3 is a full-length homolog that retains the core domain architecture of EIF4G1, including EIF4E-, EIF4A-, and EIF3-binding sites (Imataka et al. 1998). *EIF4G3* is not an essential gene (Senechal et al. 2021) and is expressed at lower levels than *EIF4G1*. However, EIF4G3 is required for spermatogenesis in the mouse (Sun et al. 2010; Senechal et al. 2021). EIF4G1 is considered the predominant scaffold for canonical cap-dependent initiation; however, the extent to which DAP5 and EIF4G3 can compensate when EIF4G1 is compromised remains an open question.

In addition to the regulation described above, quality-control pathways survey initiation and elongation to preserve translational fidelity. Colliding or stalled ribosomes are detected by surveillance factors that trigger ribosome-associated quality control (RQC). Detection of abnormal ribosome conformations results in site-specific ubiquitination of ribosomal proteins. On the small ribosomal subunit, mono-ubiquitination of RPS3 has been linked to initiation-proximal stalling (initiation RQC, iRQC) (Garshott et al. 2021), whereas modifications on RPS10 among others report elongation stalls (Meydan and Guydosh 2020). These marks interface with recycling factors, decay pathways, the proteasome, and de-ubiquitinases to resolve or clear abnormal complexes from mRNAs (Joazeiro 2019). Whether iRQC responds to loss of a core initiation scaffold, as opposed to the elongation stalls and inhibitor treatments in which it was originally characterized, has not been examined.

Defining which transcripts depend on EIF4G1, and why, requires removing the scaffold acutely. We previously developed a system to rapidly deplete EIF4G1 using an inducible degron based on the dTAG system (Nabet et al. 2020; Clark et al. 2023). Here we use this system to interrogate the consequences for translation using polysome profiling and mRNA-seq. Depletion of EIF4G1 causes a pronounced leftward shift in the polysome profile, with a loss of heavy polysomes and a large accumulation in the monosome peak, but the effect is not uniform across transcripts. Analysis of polysome-associated mRNA identifies a group of transcripts that are particularly sensitive to loss of EIF4G1, which we term Sensitive, and a second group that shows an enhancement of polysome association upon EIF4G1 depletion, which we term Resistant. mRNA feature characterization indicates that these groups correlate not only with 5′UTR features but also with open reading frame (ORF) and 3′UTR length and architecture.

Integrating our polysome profiling, RNA-seq, and reporter assays, we propose a model in which longer transcripts with GC-rich coding sequences tend to have a lower reliance on eIF4F formation, whereas shorter or low-GC transcripts rely on EIF4G1 to maintain productive initiation. In addition, we find that iRQC is activated during EIF4G1 depletion, suggesting an altered architecture of translation initiation complexes in the absence of EIF4G1. Together, these findings expand the determinants of EIF4G1 dependence beyond the 5′UTR and connect initiation surveillance to scaffold availability.

## Results

### Acute complete depletion of EIF4G1 does not affect other eIF4F components

Previously we fused a degron based on the dTAG system to the carboxy terminus of EIF4G1 using CRISPR-based methods in the HCT116 human carcinoma cell line (Nabet et al. 2020; Clark et al. 2023). In this cell line, EIF4G1 is rapidly degraded upon treatment with the small molecule dTAGV-1 (dTAG) and is no longer detectable (Figure 1A). As seen previously, despite the depletion of the EIF4G1 scaffold, levels of other components of the eIF4F complex, EIF4E and EIF4A1, remain unchanged. Additionally, the level of poly(A)-binding protein (PABPC1), another protein known to interact with EIF4G1, is unaffected by loss of the scaffold (Figure 1A).

**Figure 1.**
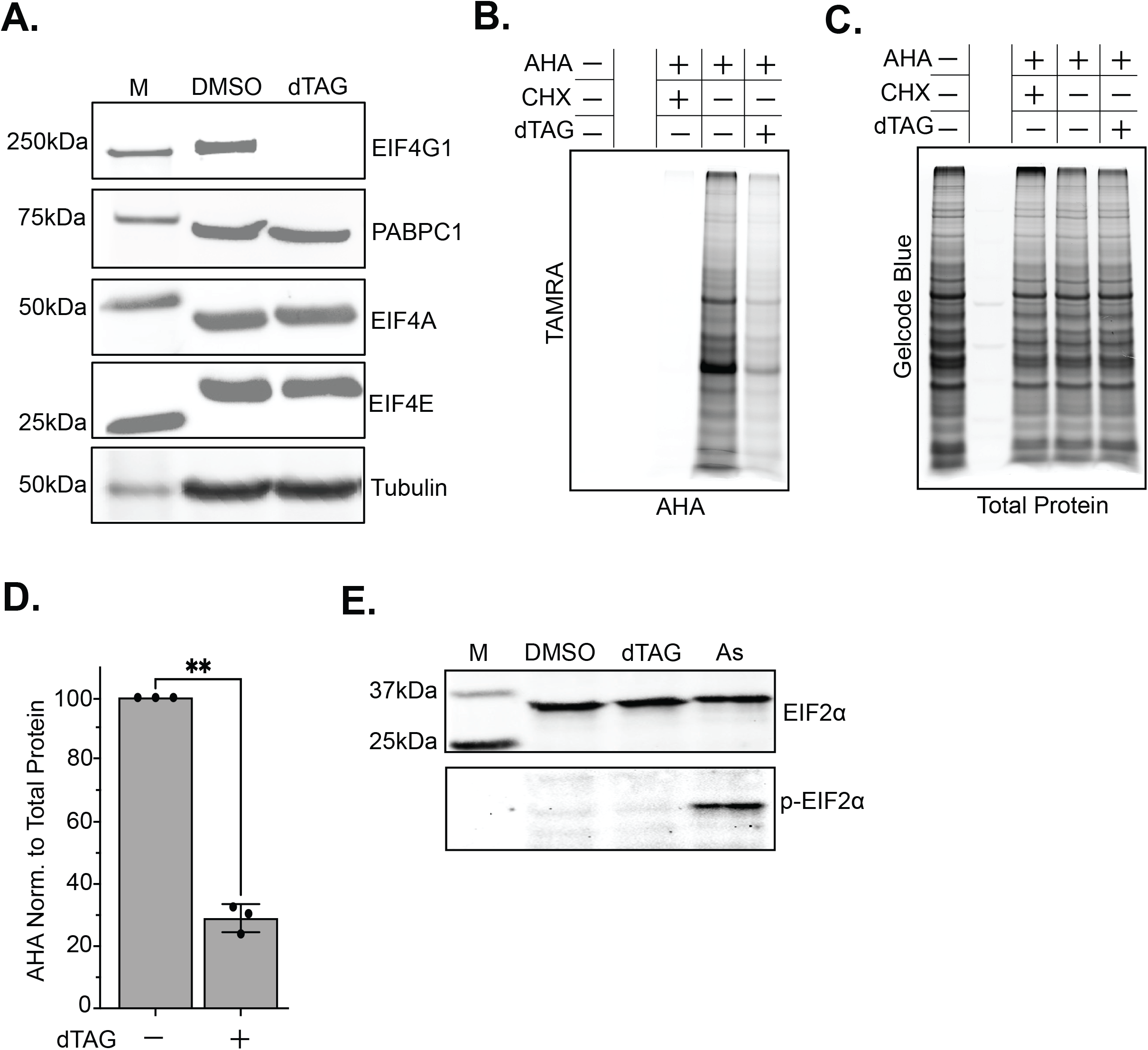
Acute EIF4G1 depletion reduces global translation without activating the integrated stress response (ISR). (A) Western blot of EIF4G1, EIF4E, EIF4A1, and PABPC1 following dTAGV-1 treatment. Tubulin was used as a loading control. (B) L-azidohomoalanine (AHA)-TAMRA labeling of nascent protein, resolved by SDS-PAGE, in cells treated with DMSO or dTAGV-1. Pretreatment with cycloheximide (CHX) was used as a negative control. (C) Coomassie staining of the gel in B as a total protein loading control. (D) Quantification of AHA incorporation in B, normalized to total protein from Coomassie staining in C (n = 3; mean ± SD). (E) Western blot of phospho-eIF2α and total eIF2α in cells treated with DMSO, dTAGV-1, or sodium arsenite (As, positive control for ISR activation).

We selected the four-hour time point, several hours after EIF4G1 becomes undetectable, which occurs within one hour of dTAG addition (Clark et al. 2023), for the analyses to determine the effects of EIF4G1 depletion on protein synthesis. This time point was chosen based on the estimated time required for ribosomes initiating via an EIF4G1-dependent mechanism to finish translation of even the longest known human transcripts. Given the elongation rate of ribosomes in mammalian cells has been measured at ∼5 codons per second (Yan et al. 1992; Ingolia et al. 2012; Wang et al. 2012), it would take approximately 2 hours to complete translation of titin which contains more than 38,000 codons (Bang et al. 2001). Any transcripts that remain associated with polysomes at this point very likely initiated translation in the absence of EIF4G1.

### Global protein synthesis is drastically impaired upon EIF4G1 depletion

To measure global translation following acute EIF4G1 loss, we used azidohomoalanine (AHA) incorporation, a methionine analog that is incorporated into nascent proteins and detected via click chemistry conjugation to a Tetramethylrhodamine (TAMRA) fluorophore (Dieterich et al. 2006) (see Materials and Methods). SDS-PAGE of lysates revealed a marked reduction in TAMRA signal in dTAG-treated cells (Figure 1B) compared to mock-treated cells, with total protein loading confirmed by Coomassie staining (Figure 1C). EIF4G1-depleted cells retained only 29% (± 4.5%) of control AHA incorporation, representing an approximately 70% reduction in global protein synthesis (Figure 1D), consistent with previous studies targeting EIF4G1 or the eIF4F complex (Stoneley et al. 2000; Sehrawat et al. 2022; Clark et al. 2023; Roiuk et al. 2024).

To determine whether this translational suppression involved activation of the integrated stress response (ISR), we monitored eIF2α phosphorylation, a hallmark of ISR-induced translational repression (Krishnamoorthy et al. 2001). Phospho-eIF2α was undetectable in both DMSO-and dTAGV-1-treated cells, while sodium arsenite, a known inducer of the ISR, produced a robust signal under identical blotting conditions (Figure 1E). Therefore, the translational defect is attributable to EIF4G1 loss rather than activation of the ISR.

### Polysome analysis reveals a redistribution of ribosomes when EIF4G1 is depleted

We separated cytoplasmic lysates on sucrose gradients to visualize the global effects on ribosome association with mRNAs. Upon EIF4G1 depletion, we observed a loss of the heaviest polysomes and a buildup of 80S monosomes (Figure 2A). To quantify the change, we determined the polysome-to-monosome (P/M) ratio, calculated as the area under the curve for the total polysome region divided by the area under the curve for the 80S monosome peak. The P/M ratio is a standard metric for quantifying global translation initiation activity from polysome profiles. Higher P/M ratios indicate more effective initiation and translation, as more ribosomes are able to efficiently load onto transcripts. The P/M ratio decreased from 2.3 ± 0.5 in DMSO-treated cells to 1.07 ± 0.12 in EIF4G1-depleted cells (Supplemental Table S1). To test whether the accumulated 80S species are vacant ribosomes rather than mRNA-associated complexes, we repeated the fractionation using 500 mM KCl, a salt concentration that dissociates vacant 80S couples (Martin and Hartwell 1970; van den Elzen et al. 2014). The redistribution to the 80S peak persisted, with no increase in free 40S or 60S subunits (Supplemental Figure S10).

**Figure 2.**
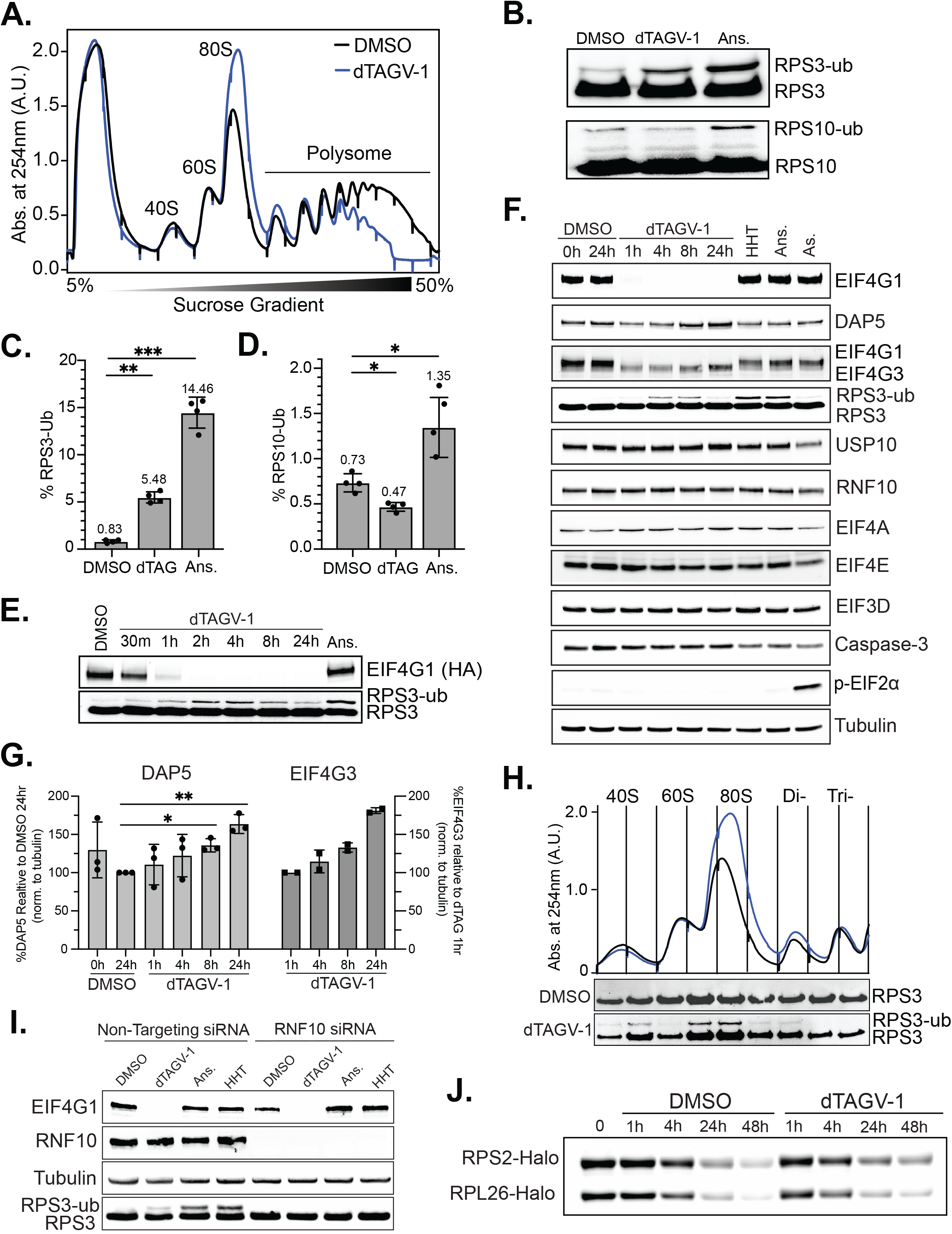
Acute EIF4G1 depletion changes the polysome profile and activates iRQC. (A) Polysome profiles (UV absorbance at 254 nm) from sucrose gradients of cells treated with DMSO or dTAGV-1. (B) Western blots of ubiquitinated RPS3 (RPS3-ub) and RPS10 (RPS10-ub) in cells treated with DMSO, dTAGV-1, or anisomycin (ANS, positive control). (C,D) Mono-ubiquitinated RPS3 (C) and RPS10 (D) as a percentage of the non-ubiquitinated form in cells treated with DMSO, dTAGV-1, or ANS (n = 4; mean ± SD). *P < 0.05, **P < 0.01, ***P < 0.001 versus DMSO, Welch’s ANOVA with Dunnett’s T3 multiple comparisons test. (E) Western blot time course of RPS3-ub following dTAGV-1 treatment. (F) Western blot time course of the indicated proteins in cells treated with DMSO (0 and 24 h), dTAGV-1 (1, 4, 8, and 24 h), homoharringtonine (HHT, 50 nM, 2 h), anisomycin (ANS, 0.1 µg/mL, 2 h), or sodium arsenite (As, 500 µM, 30 min). (G) Quantification of EIF4G2/DAP5 (n = 3; relative to the DMSO 24 h control; mean ± SD; repeated-measures one-way ANOVA with Dunnett’s multiple comparisons test) and EIF4G3 (n = 2; relative to the 1 h timepoint; individual replicate values shown) protein levels normalized to tubulin. (H) Western blot of RPS3-ub in polysome gradient fractions (40S to trisome) from cells treated with DMSO or dTAGV-1. (I) Western blots of RNF10, RPS3, and RPS3-ub following siRNA knockdown of RNF10 in cells treated with dTAGV-1, ANS, or HHT. (J) Pulse-chase of co-transfected RPS2-Halo and RPL26-Halo. Halo-TMR-labeled protein remaining at 0, 1, 4, 24, and 48 h of DMSO or dTAGV-1 treatment, measured by in-gel fluorescence.

### EIF4G1 depletion triggers initiation-linked ribosome quality control

Because EIF4G1 is thought to act before 80S assembly (Hinnebusch 2014), we expected its loss to leave 40S and 60S subunits unable to join, producing an accumulation of free subunits. Instead, neither subunit peak increased. Ribosomes redistributed into the 80S monosome peak, indicating that subunit joining still occurs when EIF4G1 is depleted. We investigated the possibility of other influences on the ribosome distribution by examining the effect of EIF4G1 depletion on the initiation-specific ribosome quality control (iRQC) pathway. To test this directly, we measured mono-ubiquitination of RPS3 (RPS3-ub) and RPS10 (RPS10-ub) (Figure 2B). Both are established indicators of RQC activation, but RPS3-ub can additionally report iRQC. Anisomycin (ANS) is an elongation inhibitor and serves as a positive control for RQC activation. ANS robustly induces both RPS3 and RPS10 mono-ubiquitination, elevating RPS3-ub by ∼17.4-fold and RPS10-ub by ∼1.84-fold relative to DMSO. Under EIF4G1 depletion, RPS3-ub increases by ∼6.6-fold (Figure 2C), whereas RPS10-ub decreases to ∼0.64-fold of DMSO (Figure 2D). This selective induction of RPS3-ub without RPS10-ub elevation is consistent with activation of iRQC rather than elongation-stage surveillance (Garshott et al. 2021).

To define the kinetics of this response, we performed a dTAGV-1 treatment time course. An increase in RPS3-ub signal relative to mock-treated cells is detectable at thirty minutes and peaks at approximately four hours after dTAG treatment (Figure 2E). The increase in RPS3-ub with a concomitant loss of RPS10-ub suggests that iRQC is the pathway activated by loss of EIF4G1.

To further address the iRQC mechanism, the loss of RPS3-ub at the later timepoint, and determine if prolonged EIF4G1 depletion activates an alternative stress response, we expanded this analysis to a dTAG treatment time course (DMSO 0 h and 24 h; dTAGV-1 1, 4, 8, and 24 h) alongside three positive controls targeting distinct quality-control and stress pathways: homoharringtonine (HHT), an initiation inhibitor used as a positive control for initiation-specific iRQC; anisomycin (ANS), an elongation inhibitor used as a positive control for general, elongation-stage RQC; and sodium arsenite (As), used as a positive control for ISR activation via eIF2α phosphorylation. We quantified known iRQC targets and RQC regulators (RPS3-ub, RNF10, USP10), initiation factors (EIF4E, EIF4A, EIF3D), an apoptosis marker (Caspase-3), and an ISR indicator (phospho-eIF2α), each normalized to tubulin (Figure 2F). EIF4E, EIF4A, EIF3D, RNF10, and USP10 showed no change in expression across the 24-hour dTAG time course. Caspase-3 levels were unchanged across all conditions tested, including the positive controls, arguing against apoptotic engagement as a confounding factor. Phospho-eIF2α was minimal throughout the DMSO and dTAG time course but was readily detected in the sodium arsenite-treated positive control, confirming that the assay is sensitive to eIF2α phosphorylation and that its absence under EIF4G1 depletion reflects a true lack of ISR activation rather than an assay limitation. Notably, EIF4G2/DAP5 levels increased over the same time course relative to the DMSO 24h control (repeated-measures one-way ANOVA with Dunnett’s multiple comparisons test), reaching statistical significance by eight hours after treatment (+35% ± 8%, n = 3; adjusted p = 0.045) and increasing further by 24 hours (+63% ± 12%, n = 3; adjusted p = 0.030; Figure 2G). Because the EIF4G3 antibody also cross-reacts with EIF4G1(Clark et al. 2023), we normalized EIF4G3 signal to the 1-hour dTAG timepoint, when EIF4G1 is no longer detectable, rather than to the DMSO 24-hour control used for EIF4G2/DAP5. Relative to this reference, EIF4G3 levels continued to increase over the remaining time course, rising by roughly 80% in each of two independent replicates by 24 hours. This finding contrasts our earlier report, where we found no change in EIF4G3 at 24 hours (Clark et al. 2023).

To pinpoint the ribosomal fraction associated with RPS3-ub, we probed sucrose gradient fractions corresponding to the 40S, 60S, and 80S ribosome, as well as the polysome region up to the trisome, for RPS3. The ubiquitinated form, RPS3-ub, is found in the 40S fraction and is enriched in the 80S monosome fraction but is absent from trisome and heavier polysome fractions (Figure 2H). This result shows that EIF4G1 depletion results in iRQC activation at the level of assembled monosomes as well as 40S-containing complexes. Recent work shows that initiation stalling can trigger iRQC, particularly under ISR activation or overexpression of the E3 ubiquitin ligase RNF10, which targets small-subunit proteins RPS2 and RPS3 (Garshott et al. 2021). Notably, neither condition is present in our system, suggesting that EIF4G1 loss alone is sufficient to engage iRQC. Together, the increase in RPS3-ub coupled with a decrease in RPS10-ub under EIF4G1 loss suggests an initiation-proximal modification that is largely independent of canonical elongation-stage RQC.

To test whether the RPS3 mono-ubiquitination we see upon EIF4G1 depletion is in the same pathway previously identified for iRQC we targeted RNF10, the E3 ligase required for canonical iRQC (Garshott et al. 2021; Garzia et al. 2021). We depleted RNF10 using siRNA and measured RPS3-ub in cells treated with DMSO, dTAGV-1, ANS, or HHT (Figure 2I). RNF10 knockdown eliminated detectable RPS3-ub under all three treatments. EIF4G1 depletion therefore engages the same RNF10-dependent ubiquitination machinery previously described for iRQC, rather than an alternative pathway.

Having observed a loss of RPS3-ub signal at the 24-hour timepoint in the dTAG time course (Figure 2F), we asked whether this reflected accelerated turnover of the ribosomal proteins themselves rather than removal of the ubiquitin mark. To test this, we generated constructs expressing HaloTag fusions of RPS2 and RPL26. The Halo protein is a modified haloalkane dehalogenase that covalently binds synthetic ligands containing a halogenated alkane (Los et al. 2008). The levels of the Halo protein can be measured in living cells through the specific labeling of the protein using commercially available functionalized chloroalkanes that contain cell-permeable fluorophores. Specific Halo labeling can be determined by separating the crude lysate by SDS-PAGE and imaging fluorescent proteins of the correct molecular weight. To confirm that these Halo-tagged ribosomal proteins are incorporated into ribosomal particles rather than existing as free, unincorporated protein, we transfected cells with RPS2-Halo and RPL26-Halo, labeled with Halo-TMR ligand, and fractionated lysates on sucrose gradients, quantifying RPS2-Halo and RPL26-Halo signal across the gradient by in-gel fluorescence. The majority of each reporter was incorporated into ribosomal subunits or fully assembled ribosomes (72.1% for RPS2-Halo, 58.9% for RPL26-Halo), with the remainder present as free, unincorporated protein (27.9% and 41.2%, respectively) (Supplemental Figure S7).

The Halo constructs were co-transfected into cells, allowing us to perform a pulse-labeling experiment. We pulse-labeled the existing pool of Halo-RPS2 and Halo-RPL26 with Halo-TMR ligand. The ligand was washed out and the cells were grown for 48 hours under DMSO or dTAG treatment. The levels of Halo-RPS2 and Halo-RPL26 were measured by quantifying the remaining TMR-labeled protein at each timepoint by in-gel fluorescence (Figure 2J). For both RPS2-Halo and RPL26-Halo, the labeled protein pool decayed steadily over the outgrowth period under both conditions, but turnover was, if anything, slower under dTAG than DMSO: by 48 hours, dTAG-treated cells retained more labeled protein than DMSO-treated cells for both RPS2-Halo (31.5% vs. 20.2% of the initial pool remaining) and RPL26-Halo (15.3% vs. 10.2% remaining). This indicates that EIF4G1 depletion does not accelerate degradation of either the small (RPS2) or large (RPL26) ribosomal subunit reporter. This experiment cannot distinguish removal of the ubiquitin mark from degradation of the modified protein, but it leaves de-ubiquitination of a persisting protein as the more likely account of the loss of RPS3-ub at later timepoints. RPS2, used here as the small-subunit reporter, is itself an RNF10 target.

### Sequencing mRNAs in heavy polysomes identifies transcripts capable of initiating in the absence of EIF4G1

Guided by the finding that iRQC is absent in tri- and higher polysomes and peaks at the four hour time point, we pooled fractions containing three or more ribosomes per transcript into a single “heavy polysome” fraction (Figure 3A) to identify transcripts that can initiate without EIF4G1 and are not bound by a ribosome containing RPS3-ub. As a proof of principle, we first used qPCR to measure candidate target-gene abundance in the heavy polysome fraction, normalized to 28S rRNA so that measurements approximate transcripts per ribosome, and calculated the log₂(dTAG/DMSO) ratio to quantify relative changes in polysome association (Figure 3B). Candidates were chosen based on published responses to eIF4F manipulation.

**Figure 3.**
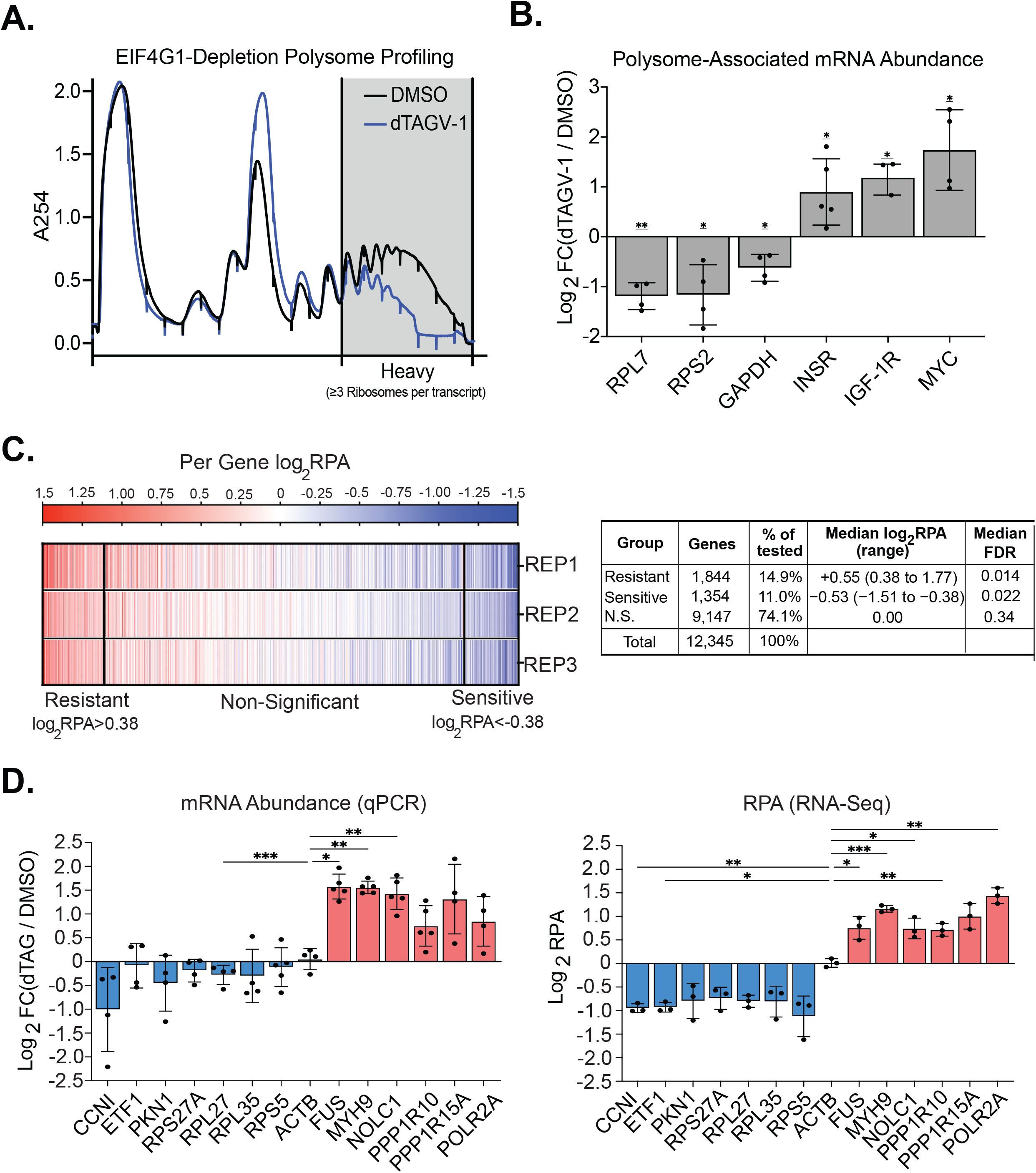
EIF4G1 depletion alters polysome association in a transcript-specific manner. (A) Polysome profiling after EIF4G1 depletion. Polysome profiles (absorbance at 254 nm) of cells treated with DMSO or dTAGV-1. Shading marks the heavy polysome fractions (≥3 ribosomes per transcript) collected for analysis. (B) Polysome-associated mRNA abundance. qPCR of candidate transcripts in the heavy polysome fraction, normalized to 28S rRNA and shown as log₂ fold change (dTAGV-1/DMSO) (n = 3 to 5; mean ± SD). *P < 0.05, **P < 0.01, one-sample t test against zero. (C) Transcriptome-wide relative polysome association (RPA). Heatmap of log₂RPA for each gene (columns) in each replicate (rows), ordered by mean log₂RPA, with summary statistics for each group. RPA is the ratio of heavy polysome to input mRNA abundance in dTAGV-1 divided by the same ratio in DMSO. Genes were classified as Resistant or Sensitive at Benjamini-Hochberg FDR < 0.05 and |log₂RPA| ≥ 0.38 (1.3-fold) using a moderated one-sample t test in limma (n = 3); all other genes are non-significant (N.S.). (D) Validation of RPA by qPCR. Log₂ fold change (dTAGV-1/DMSO) by qPCR, normalized to 28S rRNA (left; n = 4 or 5), and log₂RPA from mRNA-seq (right; n = 3) for Sensitive (blue) and Resistant (red) transcripts and ACTB (mean ± SD). *P < 0.05, **P < 0.01, ***P < 0.001 versus ACTB, Dunnett’s multiple comparisons test following a mixed-effects model (qPCR) or repeated-measures one-way ANOVA (RPA).

RPL7 and RPS2 mRNAs, which contain 5′ TOP motifs (Levy et al. 1991) and are sensitive to general inhibition of the eIF4F cap-binding complex (Thoreen et al. 2012), both showed significant losses in the heavy fraction: RPL7 showed a ∼2.3-fold decrease and RPS2 decreased ∼2.2-fold under EIF4G1 depletion. GAPDH, used as a housekeeping control previously described as relatively insensitive when targeting eIF4F complex formation (Roiuk et al. 2024), nonetheless showed an ∼1.5-fold decrease. Previously, the insulin receptor (INSR), insulin-like growth factor receptor-1 (IGF-1R), and c-myc mRNAs have been shown to contain elements within their 5′UTRs making them insensitive to eIF4F inhibition (Stoneley et al. 2000; Spriggs et al. 2009; Meng et al. 2010; Olson et al. 2013; Clark et al. 2023). In contrast to the TOP mRNAs and GAPDH, these transcripts showed increased polysome association, with INSR, IGF-1R, and c-myc increasing 1.9-fold, 2.4-fold, and 3.3-fold, respectively. Taken together, these benchmark shifts indicate real heterogeneity in EIF4G1 dependence; moreover, grouping ≥3-ribosome fractions into a combined heavy pool provides a simple readout of EIF4G1 dependence across targets, enabling reliable classification of transcripts.

### Transcriptome-wide relative polysome association identifies transcript-specific sensitivity to EIF4G1 depletion

To assess transcript-specific responses to EIF4G1 depletion, we asked how ribosome engagement shifts on individual mRNAs when initiation scaffolding is removed, while controlling for underlying mRNA abundance. We generated mRNA-seq profiles by mapping reads to GRCh38/hg38 and deriving gene-level counts from matched heavy polysome (“Heavy”) and total RNA (“Input”) samples (derived from polysome traces in Supplemental Figure S6). Batch-corrected, normalized read counts (Methods; validated by PCA, Supplemental Figure S1) were used to compute polysome association (PA = Heavy/Input) per gene, and relative polysome association (RPA = PA_dTAG_/PA_DMSO_) as the treated-to-mock ratio of PA, reported per replicate as log₂(RPA) (Figure 3C). A DESeq2 analysis of the input RNA confirmed minimal transcriptome remodeling (Supplemental Figure S2, Supplemental Table S2), indicating that changes in RPA reflect translational redistribution rather than transcript level effects. Genes were classified as

Resistant or Sensitive if their mean log₂RPA exceeded one standard deviation from the mean of the distribution in either direction (±0.38, corresponding to log₂(1.3), or approximately a 30% shift in RPA), and this shift was statistically significant by a moderated one-sample t-test (limma; (Ritchie et al. 2015)) with Benjamini-Hochberg FDR < 0.05 across all expressed genes. This identified 1,844 Resistant and 1,354 Sensitive transcripts (Supplemental Figure S3, Supplemental Table S3).

RPA measures the proportion of each transcript’s mRNA pool that remains in heavy polysomes, not its absolute ribosome occupancy. Because global protein synthesis falls by roughly 70% under these conditions, a Resistant transcript is one that retains heavy polysome association better than the average transcript, rather than one whose absolute translation increases. The analysis therefore asks which transcripts can still load ribosomes efficiently when EIF4G1 is absent.

To validate the RPA metric, we performed qPCR on 13 transcripts spanning the sensitivity spectrum, normalized to 28S rRNA, and compared each with ACTB, which showed near-zero RPA in our system (log₂RPA = 0.012). Importantly, this qPCR was performed on cDNA prepared from independent polysome gradients, not from the RNA used to generate the mRNA-seq libraries. The two measurements therefore represent separate experiments rather than two readouts of the same material. Sensitive transcripts, including CCNI, an established eIF4F-dependent mRNA (Rosenwald et al. 1993; Rousseau et al. 1996), TOP-containing ribosomal protein mRNAs (RPS5, RPL27, RPL35, RPS27A), and ETF1 and PKN1, decreased by both RPA and qPCR. Resistant transcripts, including FUS, MYH9, NOLC1, PPP1R10, PPP1R15A, and POLR2A, showed concordant increases by both methods (Figure 3D). Because the qPCR and mRNA-seq measurements were made in separate experiments, their agreement reflects reproducibility of the underlying translational shifts rather than technical concordance within a single sample set, and confirms that RPA reliably captures differential polysome association upon EIF4G1 depletion.

Analysis of individual fractions across the heavy polysome window (fractions 16 and 17; Supplemental Figure S4) showed the same direction of change: Resistant transcripts were more abundant in individual heavy fractions, whereas Sensitive transcripts, including TOP mRNAs, were reduced (Supplemental Figure S5). These patterns indicate that Resistant transcripts can maintain multi-ribosome engagement under EIF4G1 loss, whereas Sensitive mRNAs lose heavy-fraction association consistent with impaired ribosome loading.

### Sequence features associated with transcript classes

In an effort to determine if the mRNAs in each of the identified classes (Resistant and Sensitive) have any related functions we used a gene ontology enrichment analysis. This approach revealed that Resistant and Sensitive transcripts are enriched for several distinct biological functions (Supplemental Figure S9). Resistant transcripts were enriched for transcriptional regulators as well as regulators of cell migration. Sensitive transcripts were most strongly enriched for the ontology group defined as cytoplasmic translation (6.4-fold), including the small and large ribosomal subunits (6.9- and 6.0-fold). Many of these transcripts are canonical 5′ terminal oligopyrimidine (TOP) mRNAs, consistent with a TOP motif enrichment in this group.

This finding that a motif was associated with one class of transcripts raised the question of whether specific sequence or structural features within these transcripts drive their differential sensitivity to EIF4G1 depletion. An alternative to this is that one of the other EIF4G family members is driving the classification. EIF4G2 (DAP5/NAT1) is a homolog of EIF4G1 that shares its central HEAT domain and lacks the EIF4E-binding region but retains the ability to recruit eIF4A (Imataka et al. 1997). *EIF4G2* is also highly expressed in this cell line while *EIF4G3* is expressed at only low levels. To ask whether EIF4G2 might contribute to the continued translation of some transcripts when EIF4G1 is absent, we compiled a list of DAP5-dependent mRNAs from three published studies (Yoffe et al. 2016; David et al. 2022; Weber et al. 2022), identifying 467 unique genes, of which 422 passed the expression thresholds required for classification in our dataset (313 Non-significant, 44 Sensitive, 65 Resistant). DAP5-dependent transcripts were not over-represented among Resistant transcripts: 65 of 422 (15.4%) were classified Resistant and 44 (10.4%) Sensitive, closely matching the proportions across all classified genes (14.9% and 11.0%; chi-square test of independence, chi-square = 0.18, df = 2, P = 0.91). DAP5 dependence therefore does not predict resistance to EIF4G1 loss.

We examined sequence features associated with sensitivity or resistance upon EIF4G1 depletion and included this DAP5-dependent group as a fourth comparison group alongside our Non-significant, Sensitive, and Resistant classifications in the sequence-feature analyses below across all four groups. We first examined transcript length (Figure 4A), partitioning into 5′UTR, CDS, and 3′UTR, as well as total transcript length. Transcript length was significantly associated with the effects of EIF4G1 depletion: Sensitive transcripts tended to be shorter overall and across 5′UTR, CDS, and 3′UTR, whereas Resistant transcripts tended to be longer in each region. The DAP5-dependent set also trended toward longer transcripts, particularly in the 5′UTR, consistent with the known role of DAP5 in resolving structured 5′UTR elements (Weber et al. 2022); Resistant transcripts share this trend toward longer 5′UTRs. It is important to note that these are population-level trends with substantial overlap between distributions; transcript length is not a binary determinant of EIF4G1 dependence. The CDS showed the most pronounced length difference among the Non-significant, Sensitive, and Resistant groups, consistent with findings in yeast where eIF4G depletion preferentially affects translation of shorter mRNAs (Park et al. 2011).

**Figure 4.**
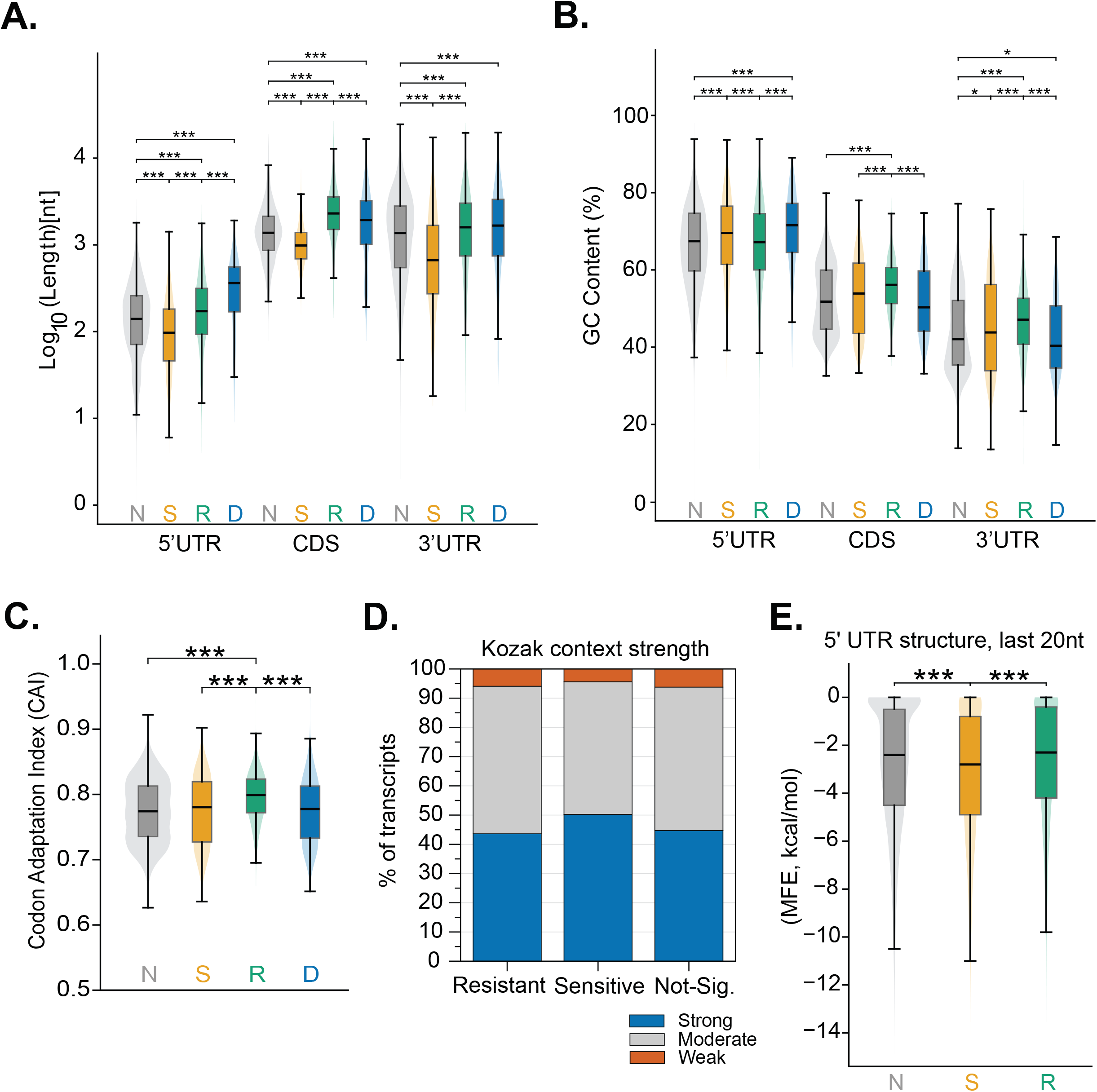
Sequence features associated with EIF4G1 dependence and comparison with DAP5-dependent transcripts. (A) Length distributions of the 5′UTR, CDS, and 3′UTR for Non-significant (N), Sensitive (S), Resistant (R), and DAP5-Dependent (D) transcripts. The DAP5-Dependent set (n = 422) is drawn from published studies, and its members also belong to one of the other three groups. (B) GC content distributions of the same regions for the same four groups. (C) Codon adaptation index (CAI) distributions for the same four groups. (D) Percentage of transcripts with strong, moderate, or weak Kozak context in the N, S, and R groups. (E) Minimum free energy (MFE; kcal/mol) of predicted secondary structure in the terminal 20 nt of the 5′UTR for the N, S, and R groups. In A, B, C, and E, violin plots contain embedded box plots, and P values are from pairwise Mann-Whitney U tests.

GC content analysis revealed a region-specific pattern (Figure 4B): Resistant transcripts had significantly higher GC content in the CDS and 3′UTR, while DAP5-dependent transcripts trended below the genome-wide median in CDS GC content, distinguishing the two groups despite their shared trend toward longer 5′UTRs. The 5′UTR showed a different trend among the Non-significant, Sensitive, and Resistant groups: Sensitive transcripts had higher 5′UTR GC content than Resistant transcripts. This suggests that GC content may operate under distinct functional constraints in the 5′UTR compared to the coding region: higher 5′UTR GC content may increase secondary structure and scanning barriers, whereas higher CDS GC content may be associated with reduced eIF4F dependence through alternative mechanisms. As with transcript length, these are distributional trends with overlap between groups, and GC content alone does not predict EIF4G1 dependence; the divergent CDS GC content of the Resistant and DAP5-Dependent groups further suggests that the features associated with translational resistance upon EIF4G1 loss are distinct from those associated with DAP5-driven initiation. To ask whether specific regulatory elements are associated with EIF4G1 dependence independent of bulk sequence features, we examined a curated set of established 5′ terminal oligopyrimidine (TOP) mRNAs (Supplemental Table S7). Of 92 TOP mRNAs, 60 were Sensitive, a 5.95-fold enrichment over the Sensitive fraction of all transcripts (p = 1.34 x 10^-35^), consistent with the established eIF4F-dependence of TOP mRNAs (Thoreen et al. 2012; Jia et al. 2021).

### Higher CDS GC content promotes translation under EIF4G1 depletion in reporter assays

Among the structural features examined, CDS GC content stood out as a feature that differentiated Resistant transcripts (Figure 4B) and had not been identified in previous analysis of EIF4G usage. Codon adaptation index (CAI) was also significantly different between groups, with Resistant transcripts showing higher median CAI than Non-significant, Sensitive, or DAP5-

Dependent transcripts (Resistant median CAI = 0.799 vs. 0.774 to 0.781 across the other three groups; Mann-Whitney p < 10^-18^ for every comparison against Resistant; Supplemental Table S8). However, the magnitude of this separation was substantially smaller than for CDS GC content itself: CAI increased by only ∼3% relative to Non-significant (+0.025), compared to an ∼8% relative increase in CDS GC% over the same comparison (+4.3 percentage points), indicating that GC content within the CDS, rather than codon usage bias, is a dominant compositional signature of the Resistant class (Figure 4C).

We next examined the Kozak translation initiation context (a purine at position −3 and a guanine at position +4 relative to the start codon) across classification groups (Figure 4D). Sensitive transcripts showed a significantly higher proportion of strong Kozak contexts than either Non-significant or Resistant transcripts (52.4% vs. 46.2% and 44.7%, respectively; p = 1.5 x 10^-5^ and p = 3.4 x 10^-4^), while Resistant did not differ significantly from Non-significant.

To ask whether local RNA secondary structure near the start codon differs by classification group, we computed the minimum free energy (MFE) of the terminal 20 nucleotides of the 5′UTR (RNAfold; Figure 4E). Sensitive transcripts showed significantly more stable local structure than both Non-significant and Resistant transcripts (median MFE = −2.8 vs. −2.4 and −2.3 kcal/mol; p = 5.2 x 10^-5^ and p = 8.1 x 10^-6^), while Resistant did not differ from Non-significant (p = 0.11). This is consistent with the elevated 5′UTR GC content already noted among Sensitive transcripts (Figure 4B).

Given that the relationship of CDS GC content to EIF4G usage has not been observed previously, we sought to validate this finding in a reporter based assay. To test whether CDS GC content can influence translation during EIF4G1 depletion, we first engineered two self-labeling Halo reporter constructs with varied GC content but identical amino acid sequences: Low-GC (43% GC, CAI = 0.74) and High-GC (65% GC, CAI = 0.84), driven by a CMV promoter and containing the same adenovirus tripartite leader (TPL) 5′UTR (Logan and Shenk 1984), which strongly reduces the requirement for eIF4F, and the same 3′UTR. Using a single 5′ and 3′UTR across all constructs holds the UTRs constant, so that differences between constructs are attributable to the CDS (Figure 5A).

**Figure 5.**
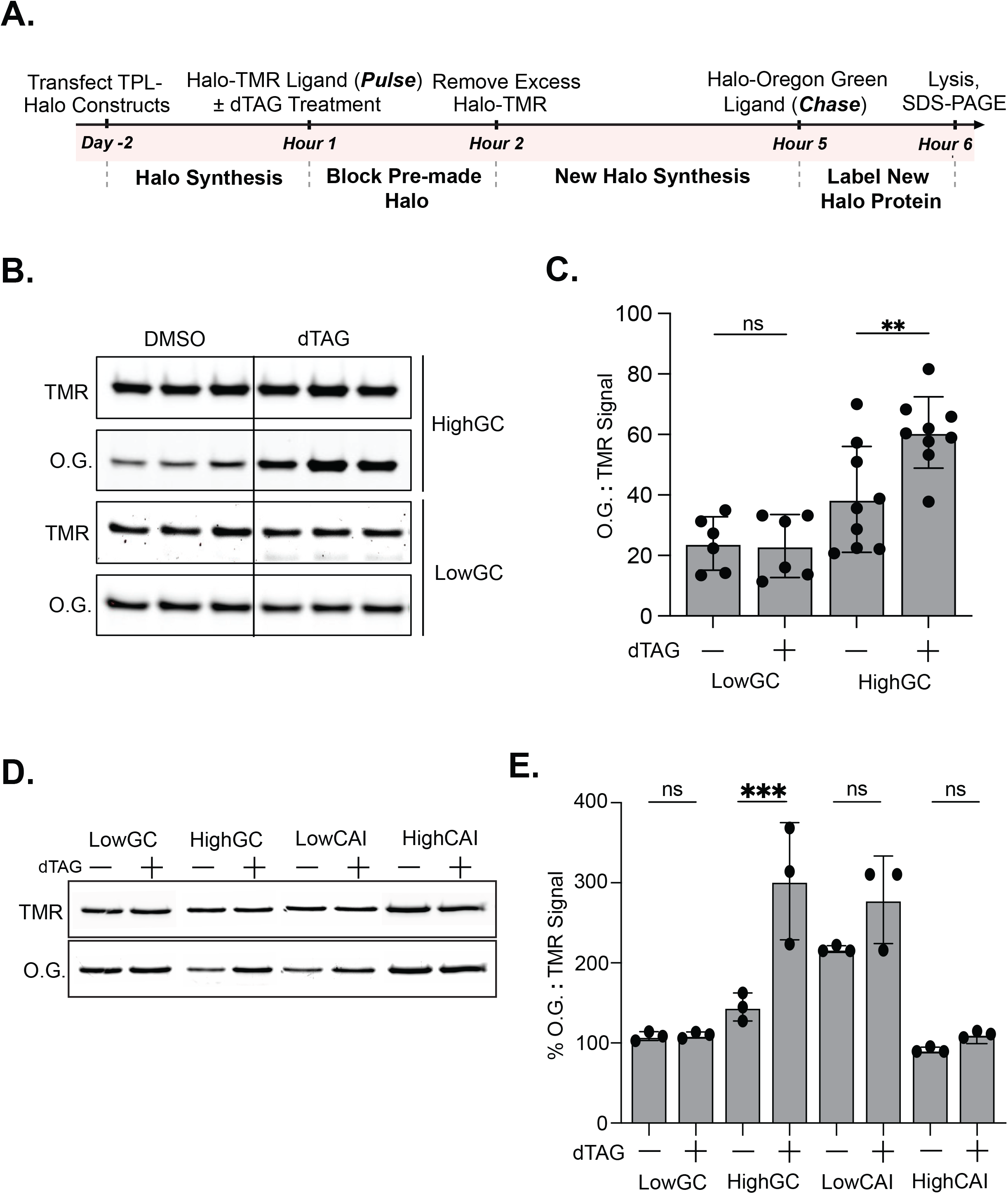
CDS GC content modulates translation during EIF4G1 depletion in a reporter assay. (A) Schematic of the Halo pulse-chase reporter assay. (B) Representative fluorescent SDS-PAGE gel of Low-GC and High-GC reporters from cells treated with DMSO or dTAGV-1, showing TMR (preexisting protein) and Oregon Green (newly synthesized protein) signals. (C) Quantification of the Oregon Green:TMR ratio for the reporters in B (Low-GC n = 6, High-GC n = 9; mean ± SD; ordinary one-way ANOVA with Tukey’s multiple comparisons test). ns, Not significant; **P < 0.01. (D) Representative fluorescent SDS-PAGE gel, as in B, of CAI-matched Low-GC and High-GC reporters and GC-matched Low-CAI and High-CAI reporters. (E) Quantification of the Oregon Green:TMR ratio for the reporters in D (n = 3; mean ± SD; one-way ANOVA with Tukey’s multiple comparisons test).

Comparing DMSO and dTAG treatment for each construct, the Low-GC reporter showed no change in signal, while the High-GC reporter showed significantly increased translation upon EIF4G1 depletion (p = 0.0074, Tukey’s following one-way ANOVA) (Figure 5B, C). However, because GC content and codon optimality co-vary in these constructs, this result alone could not distinguish whether GC content or codon usage was responsible for the increase. To decouple these two features, we engineered two additional pairs of Halo reporter constructs, each encoding an identical HaloTag amino acid sequence but differing only in synonymous codon usage: a GC-varying, CAI-matched pair, Low-GC (41% GC) and High-GC (65% GC), both held at CAI of approximately 0.82 to 0.83, isolating the effect of GC content independent of codon optimality; and a GC-matched, CAI-varying pair, Low-CAI (CAI = 0.50) and High-CAI (CAI = 0.93), both held at approximately 50% GC, isolating the effect of codon optimality independent of GC content. All four constructs maintained a minimum per-codon weight above 0.25, so no rare codons were introduced in any construct, and all shared the same CMV promoter, TPL 5′UTR, and 3′UTR, so that only CDS composition varied across constructs (Supplemental Table S9). Cells transfected with each construct were pulsed with Halo-TMR to saturate and block the preexisting protein pool, then chased with Halo-Oregon Green to selectively label protein synthesized during EIF4G1 depletion. Comparing DMSO and dTAG treatment within each construct (Tukey’s multiple comparisons test), only the High-GC construct showed a significant change in Halo signal, with output approximately doubling upon EIF4G1 depletion (Figure 5D, E; p = 0.0005). The Low-GC, Low-CAI, and High-CAI constructs each showed no significant change between DMSO and dTAG conditions (p > 0.9999, p = 0.3617, and p = 0.9947, respectively), including no effect for High-CAI despite its large CAI difference from Low-CAI at matched GC content. This indicates that CDS GC content, not codon optimality, is sufficient to confer a translational advantage upon EIF4G1 depletion. Biological replicates 2 and 3 showed consistent results (Supplemental Figure S8).

## Discussion

The system described here enables rapid and selective degradation of EIF4G1 without destabilizing other core translation initiation factors associated with eIF4F, allowing us to capture a snapshot of translation following the loss of the canonical eIF4F scaffold. Because EIF4E, EIF4A1, and PABPC1 remain intact, the observed translational defects can be attributed to EIF4G1 loss rather than a loss of the eIF4F initiation machinery. This acute perturbation is designed to isolate primary initiation defects from slower adaptive responses and provides a defined temporal window for mechanistic analysis.

Acute EIF4G1 depletion causes heavy polysome collapse with accumulation of 80S monosomes. The accumulation of 80S ribosomes without a rise in free 40S or 60S subunits, alongside a profound reduction in nascent protein synthesis, indicates that ribosomes continue to assemble on mRNA but fail to produce protein, rather than failing to be recruited at all. This phenotype is consistent with engagement of iRQC, in which RNF10-mediated mono-ubiquitination of RPS3 marks initiation-arrested complexes (Garshott et al. 2021). The enrichment of RPS3-ub in 40S and 80S species, but its absence from trisome and heavier polysomes, supports a model in which iRQC acts at or near start-codon commitment, marking ribosomes that stall before productive elongation. In this framework, EIF4G1 loss may convert the translational landscape into one characterized by poorly productive initiation intermediates, which could explain how amino acid incorporation drops precipitously while ribosomes remain assembled on mRNA.

Despite the ∼6.6-fold increase in RPS3-ub upon EIF4G1 loss, the modified form represents only ∼5% of total RPS3. However, this likely underestimates the true extent of iRQC engagement, as prior work has shown that the E3 ubiquitin ligase RNF10 is limiting relative to the de-ubiquitinase activity that opposes it (Garshott et al. 2021), meaning RPS3-ub marks are rapidly cleared and the signal we detect likely represents only a fraction of the initiation-stalled events occurring upon EIF4G1 depletion. The observation that EIF4G1 loss triggers iRQC in the absence of ISR activation suggests that scaffold depletion alone is sufficient to engage initiation-specific quality control, independent of the canonical phospho-eIF2α pathway.

Our transcriptome-wide RPA analysis reveals trends suggesting that EIF4G1 dependence is shaped by whole-transcript architecture rather than 5′UTR properties alone. Resistant transcripts tend to be longer and the most GC-rich in the CDS and 3′UTR, whereas Sensitive transcripts tend to be shorter and intermediate in GC content in these regions, consistent with findings in yeast where depletion of the EIF4G1 ortholog TIF4631 disproportionately reduced translation of shorter mRNAs (Park et al. 2011). However, it is important to emphasize that these are distributional profiles with substantial overlap between the characteristics of the classes. Many individual transcripts do not conform to these population-level trends, and transcript length or GC content alone is not sufficient to predict EIF4G1 dependence for a given mRNA.

Analysis of established TOP mRNAs reinforces this model: curated TOP mRNAs are 5.95-fold enriched among Sensitive transcripts, recapitulating the established mTORC1-LARP1-eIF4F regulatory axis (Levy et al. 1991; Thoreen et al. 2012). Sensitive transcripts also show higher 5′UTR GC content than Resistant transcripts. Whether this reflects greater secondary structure cannot be assessed across the full 5′UTR: predicted folding energy scales with sequence length, and 5′UTR length differs systematically between groups (Figure 4A), so full-length MFE values cannot be normalized for comparison between groups. We therefore examined a fixed-length window, the terminal 20 nucleotides of the 5′UTR immediately upstream of the start codon, which removes length as a variable. We focused on this region because transcriptome-wide structure probing under eIF4A inhibition with hippuristanol found that eIF4A-dependent mRNAs are distinguished by increased structure immediately upstream of the start codon (Waldron et al. 2019). Sensitive transcripts show significantly more stable predicted structure in this window than either Resistant or Non-significant transcripts (Figure 4E). This fits the canonical model of eIF4F-dependent initiation: EIF4G1 loss impairs delivery of eIF4A to the 5′UTR, and transcripts with structured elements near the start codon, which depend most on eIF4A-mediated unwinding for the scanning ribosome to reach the AUG, are the ones most compromised when EIF4G1 is lost.

If an alternative scaffold sustains initiation on Resistant transcripts after EIF4G1 loss, EIF4G2 (DAP5) is the most plausible candidate. It is the most abundant eIF4G paralog after EIF4G1: in HCT116 proteomes, EIF4G2 protein is present at roughly half the level of EIF4G1, whereas EIF4G3 is more than an order of magnitude lower (Gholami et al. 2013; Frejno et al. 2017) (Supplemental Table S10). In contrast, EIF4G2 mRNA is approximately five-fold more abundant than EIF4G1 mRNA in our HCT116 RNA-seq (Supplemental Table S10), implying roughly 9-fold lower protein output per mRNA. If DAP5 and EIF4G1 served distinct sets of mRNAs, transcripts that depend on DAP5 should be enriched among those that tolerate EIF4G1 loss. They are not. Transcripts identified as DAP5-dependent in genome-wide studies (Yoffe et al. 2016; David et al. 2022; Weber et al. 2022) were distributed across the Sensitive, Resistant, and Non-significant classes in the same proportions as all transcripts. Moreover, in the 5′UTR, the region most directly involved in initiation, DAP5-dependent transcripts share the elevated GC content of Sensitive transcripts rather than the lower 5′UTR GC content of Resistant transcripts, and they lack the high CDS GC content that distinguishes the Resistant class (Figure 4B). Because DAP5-dependent regulation is thought to act through the 5′UTR, that region is the most informative for this comparison. Together, these observations suggest that DAP5 and EIF4G1 favor overlapping rather than distinct mRNA populations. A mechanistic basis for this overlap has been reported: although DAP5 cannot bind the cap, its recruitment to target mRNAs requires the cap, eIF4F, and eIF4A, and DAP5 acts after cap-dependent ribosome loading to promote translation of the main coding sequence (Weber et al. 2022). DAP5 therefore appears to work alongside EIF4G1-containing eIF4F rather than as an independent route around it, and DAP5 dependence does not explain why Resistant transcripts tolerate EIF4G1 loss.

Our transcriptome-wide analysis points to CDS GC content as the strongest compositional predictor of EIF4G1 dependence, differing significantly between Resistant transcripts and every other group. Resistant transcripts also have modestly higher codon adaptation index (CAI) than Sensitive transcripts, raising the possibility that codon optimality, rather than GC content itself, underlies this difference. Codon optimality and GC content are naturally correlated across the transcriptome as human cells prefer GC rich codons. However, our reporter experiments, using two independent construct pairs, one of which explicitly decoupled GC content from CAI, show that GC content, not codon usage, is the feature sufficient to confer a translational advantage upon EIF4G1 depletion. Both reporter pairs share the TPL 5′UTR, which minimizes 5′UTR-driven differences in eIF4F dependence between constructs. Because the Low-GC and High-GC constructs are CAI-matched and share this 5′UTR, a model in which EIF4G1 dependence is set by the 5′UTR alone predicts no difference between them upon EIF4G1 depletion. Instead, the High-GC construct outperforms the Low-GC construct. This leaves two potential possibilities: under EIF4G1 depletion, the High-GC construct, which shares the same loading mechanism as Low-GC, either elongates more efficiently, or the GC rich transcript localizes to ribosomal subunits more effectively.

Together, our findings establish that EIF4G1 dependence is shaped by transcript-wide architectural features, including CDS length and GC content, rather than 5′UTR properties alone, and that acute loss of the canonical initiation scaffold is sufficient to activate iRQC without ISR induction, indicating that reduced eIF2*a* availability does not account for the observed ribosome quality control response. Whether these two outcomes are mechanistically linked remains to be determined; it is possible that transcripts unable to sustain productive initiation without EIF4G1 accumulate as nonproductive 80S complexes subject to RPS3 ubiquitination. Future work defining the relationship between transcript architecture, translational sensitivity, and initiation-specific quality control will be important for understanding how cells surveil and respond to disruptions in the initiation machinery.

## Materials and Methods

### Cell culture and drug treatment

Human HCT116 cells harboring FKBP12(F36V)-tagged EIF4G1 were cultured in DMEM (high glucose) supplemented with L-glutamine, sodium pyruvate, 10% FBS, and 1% penicillin-streptomycin. Cells were passaged using Trypsin-EDTA (0.25%). For EIF4G1 depletion, cells were treated with 1 µM dTAGV-1 (Tocris) for 4 hours unless otherwise noted. Anisomycin (Ans.) was used at 0.1 µg/mL for 2 hours as a positive control for RQC activation. Homoharringtonine (HHT) was used at 50 nM for 2 hours. Sodium arsenite (As.) was used at 500 µM for 30 minutes as a positive control for ISR activation. Vehicle controls received matched DMSO.

Plasmid transfections were performed using Lipofectamine 3000 (ThermoFisher) according to the manufacturer’s protocol.

### siRNA knockdown

RNF10 was depleted using ON-TARGETplus SMARTpool siRNA targeting human RNF10 (Dharmacon, L-006918-00-0005), with ON-TARGETplus Non-targeting Pool (Dharmacon, D-001810-10-05) as a control. siRNAs were delivered at 50 nM final concentration with Lipofectamine RNAiMAX (ThermoFisher) according to the manufacturer’s protocol, in two sequential transfections on consecutive days. Cells were harvested 72 hours after the second transfection, with dTAGV-1 (1 µM) added for the final 4 hours and anisomycin (0.1 µg/mL) or homoharringtonine (50 nM) for the final 2 hours. Knockdown was confirmed by western blot for RNF10 (Figure 2I).

### AHA incorporation assay

Cells were cultured in methionine-free medium for 2 hours, then treated with DMSO or 1 µM dTAGV-1 for 2 hours. Cells were subsequently labeled with 50 µM AHA·HCl (Tocris, 6584) for 2 hours (± dTAG). Lysates were prepared in RIPA buffer with Pierce Universal Nuclease and subjected to click chemistry with 1mM CuSO_4_, 5mM THPTA, 100µM tetramethylrhodamine-alkyne (ThermoFisher, T10183), and fresh 2.5mM sodium ascorbate. Proteins were methanol-precipitated, dried on ice, and solubilized in 6M urea (containing 50mM Tris pH 7.5 and 1% SDS), resolved on 8% Bis-Tris gels, and imaged for TAMRA fluorescence followed by Coomassie staining for total protein normalization.

### Western blotting and mono-ubiquitination quantification

Samples were denatured in LDS with 100 mM DTT and resolved on Bis-Tris gels in MOPS running buffer. Proteins were transferred to PVDF membranes (wet transfer, 90 V, 2 hours). Membranes were blocked in 5% non-fat dry milk/TBST and probed with indicated primary and secondary antibodies (Supplemental Table S4). For mono-ubiquitination quantification, RPS3-ub and RPS10-ub band intensities were background-subtracted and normalized to total RPS3 or RPS10, respectively. Replicates were summarized as mean ± SD; values were further normalized to DMSO for between-blot comparison.

### Sucrose gradient fractionation and polysome profiling

Step gradients (5–50% sucrose) were prepared in gradient buffer (20 mM HEPES pH 7.6, 100 mM KCl, 5 mM MgCl₂, 100 µg/mL cycloheximide). After dTAG treatment, ribosomes were trapped with cycloheximide (100 µg/mL, 15 min, 37°C). Cells were lysed in hypotonic buffer supplemented with Triton X-100 and sodium deoxycholate. Clarified lysates (10–15 A₂₆₀ units) were loaded onto gradients and centrifuged at 35,000 rpm in an SW41 Ti rotor for 2 hours at 4°C. Gradients were fractionated at 1.5 mL/min into 500 µL fractions. For high-salt gradients, KCl in the gradient and lysis buffers was raised to 500 mM, with all other conditions unchanged. Polysome-to-monosome (P/M) ratios were calculated from the area under the curve (baseline-subtracted) for the 80S monosome peak and for the total polysome region of each trace, quantified in GraphPad Prism 11 (Supplemental Table S1).

### RPS2-Halo and RPL26-Halo pulse-chase and incorporation assays

Cells were transfected with RPS2-Halo or RPL26-Halo plasmids under a CMV promoter. For the pulse-chase decay assay (Figure 2J), the existing protein pool was labeled with HaloTag-TMR ligand (1:1000, 1 hour), then cells were lifted from the plate and washed three times in PBS before DMSO or dTAGV-1 treatment was followed over the indicated time course. For the ribosomal incorporation assay (Supplemental Figure S7), cells were transfected for 2 days with no DMSO/dTAG treatment, labeled with HaloTag-TMR ligand as above, then flash-frozen, lysed, and fractionated on sucrose gradients as described above. Labeled protein was visualized by SDS-PAGE and in-gel fluorescence imaging (Bio-Rad imager).

### RNA purification, library preparation, and sequencing

We chose polysome profiling coupled with mRNA-seq over ribosome profiling to enable transcript-wide quantification of ribosome association across 5′UTR, CDS, and 3′UTR regions. RNA was extracted from total cell lysates or pooled heavy polysome fractions (≥3 ribosomes) using TRIzol/chloroform extraction with glycogen carrier and ethanol or isopropanol precipitation. Libraries were prepared using the NEBNext rRNA Depletion Kit v2 (NEB E7405) followed by the Illumina TruSeq RNA Sample kit. Libraries were quantified on the Agilent D1000 ScreenTape system.

### RNA-seq preprocessing, normalization, and batch correction

Paired-end FASTQ files were preprocessed with fastp (Phred ≥ 20, minimum insert ≥ 30 nt). Trimmed reads were aligned with STAR to GRCh38 using GENCODE v47 gene models. Raw gene-level counts were filtered (summed counts ≥ 120 per fraction, no zeros), normalized by DESeq2 size factors (Love et al. 2014), log₂-transformed (pseudocount 0.5), batch-corrected with ComBat (Johnson et al. 2007), and back-transformed. Batch correction was validated by principal component analysis on raw, normalized, and ComBat-corrected counts (Supplemental Figure S1). Mapping Metrics can be found in Supplemental Table S5.

### RPA calculation and gene classification

Polysome association (PA = Heavy/Input) was computed on normalized, batch-corrected values for DMSO and dTAG separately. RPA = PA_dTAG_/PA_DMSO_, reported as log₂(RPA) per replicate. Genes were classified as Resistant or Sensitive if (1) their mean log₂RPA exceeded one standard deviation from the mean of the distribution in either direction (±0.38, corresponding to log₂(1.3), or approximately a 30% shift in RPA), and (2) this shift was statistically significant by a moderated one-sample t-test implemented in limma (Ritchie et al. 2015), which uses empirical Bayes variance shrinkage to stabilize per-gene variance estimates across replicates, followed by Benjamini-Hochberg correction across all 12,345 expressed genes (BH-FDR < 0.05).

### Dominant transcript selection

For each gene, a single dominant transcript was selected using a two-tier, annotation-based scheme independent of expression in this dataset. Tier 1 used the MANE Select transcript (the NCBI RefSeq/Ensembl-GENCODE consensus reference isoform) where annotated (19,225 genes). Tier 2, for genes without a MANE Select transcript, used the APPRIS principal isoform (GENCODE/APPRIS annotation based on cross-species conservation and protein structural evidence), preferring the highest-confidence principal tier and breaking residual ties by transcript coverage (851 genes; principal_1: 692, principal_2: 24, principal_3: 1, principal_4: 7, principal_5: 127). Three genes lacking both annotations were excluded. This yielded one dominant transcript for 20,076 genes (GENCODE v47, 2024-07-19), of which 12,345 had a limma-based RPA classification.

### Sequence feature analysis

GC content was calculated per transcript region (5′UTR, CDS, 3′UTR) directly from the dominant transcript sequence. Codon Adaptation Index (CAI) was computed as the geometric mean of per-codon relative synonymous codon usage weights (Sharp and Li 1987); the same reference weight table was used for the transcriptome-wide analysis and the four Halo reporter constructs. Kozak context strength was classified using established consensus criteria (Kozak 1986): a purine at position −3 and a guanine at position +4 were each scored present or absent, with both present classified as strong, one as moderate, and neither as weak. Local secondary structure of the terminal 20 nucleotides of the 5′UTR was predicted by minimum free energy (MFE) folding using RNAfold (ViennaRNA package). Established 5′ terminal oligopyrimidine (TOP) mRNAs were defined by a curated list compiled from published sources (Supplemental Table S7) (Levy et al. 1991).

### RT-qPCR validation

RNA for qPCR was obtained from polysome gradients prepared independently of those used for mRNA-seq library preparation, so the qPCR and sequencing datasets derive from separate biological replicates. 5 µg RNA was reverse transcribed with oligo(dT) and random hexamers using MMLV reverse transcriptase. qPCR was performed with SYBR Green detection. Target primers were sourced from PrimerBank or designed with Primer3 (Supplemental Table S6). For heavy-fraction qPCR, candidate gene abundance was normalized to 28S rRNA. For RPA validation, candidate gene abundance was normalized to 28S rRNA, and ACTB was included as an unchanged reference transcript.

### Halo GC reporter assay

Plasmids containing Halo reporter constructs (Supplemental Table S9) were transfected and cultured for 54 hours, pretreated with DMSO or dTAGV-1 (1 hour), then pulse-labeled with 5 µM HaloTag-TMR (1 hour) to block preexisting protein. After washing, cells were returned to medium ± dTAGV-1 for 3 hours, then chased with 1 µM HaloTag-Oregon Green (1 hour). Lysates were resolved by SDS-PAGE and imaged for TMR and Oregon Green fluorescence. Translation output was quantified as Oregon Green:TMR ratio.

### DAP5-dependent transcript comparison

DAP5-dependent genes were compiled from three published genome-wide studies (Yoffe et al. 2016; David et al. 2022; Weber et al. 2022), yielding 467 unique genes. Gene symbols were mapped to current HGNC nomenclature where aliases had changed. Of these, 422 passed the expression thresholds required for classification in our dataset. Structural feature comparisons between DAP5-dependent transcripts and our three EIF4G1 classification groups used Mann-Whitney U tests. Enrichment of DAP5-dependent genes across the EIF4G1 classification groups was tested by chi-square test of independence, comparing the distribution of DAP5-dependent genes across the Resistant, Sensitive, and Non-significant classes with that of all other classified genes.

### Statistical analysis

All statistical tests were performed in Python using scipy.stats, except where noted. Fisher’s exact tests were used for overlap enrichment analyses. Mann-Whitney U tests were used for continuous feature comparisons between groups. Multiple testing correction used the Benjamini-Hochberg procedure. Ribosomal protein ubiquitination (Figures 2C and 2D) was compared across treatments by Welch’s ANOVA with Dunnett’s T3 multiple comparisons test. Halo reporter Oregon Green:TMR ratios (Figures 5C and 5E) were compared by ordinary one-way ANOVA with Tukey’s multiple comparisons test. For heavy fraction qPCR (Figure 3B), log₂ fold changes were compared with zero using two-tailed one-sample t tests. For RPA validation (Figure 3D), each transcript was compared with ACTB using Dunnett’s multiple comparisons test following a mixed-effects model (qPCR) or repeated-measures one-way ANOVA (log₂RPA), with the Geisser-Greenhouse correction (GraphPad Prism). DAP5 protein levels across the dTAG time course were compared to the DMSO 24h control using a repeated-measures one-way ANOVA with Dunnett’s multiple comparisons test (GraphPad Prism).

## Acknowledgments

[Acknowledgments and funding sources to be added.]

## Author Contributions

[Z.D.K. and M.T.M. designed the experiments. Z.D.K. performed the experiments and analyzed the data. Z.D.K. and M.T.M. wrote the manuscript.]

## Data Availability

[RNA-seq data have been deposited in the NCBI Gene Expression Omnibus (GEO) under accession number GSE######. All other data supporting the findings of this study are available within the paper and its supplemental materials.]

## Supplemental Figure Legends

**Figure. S1.**
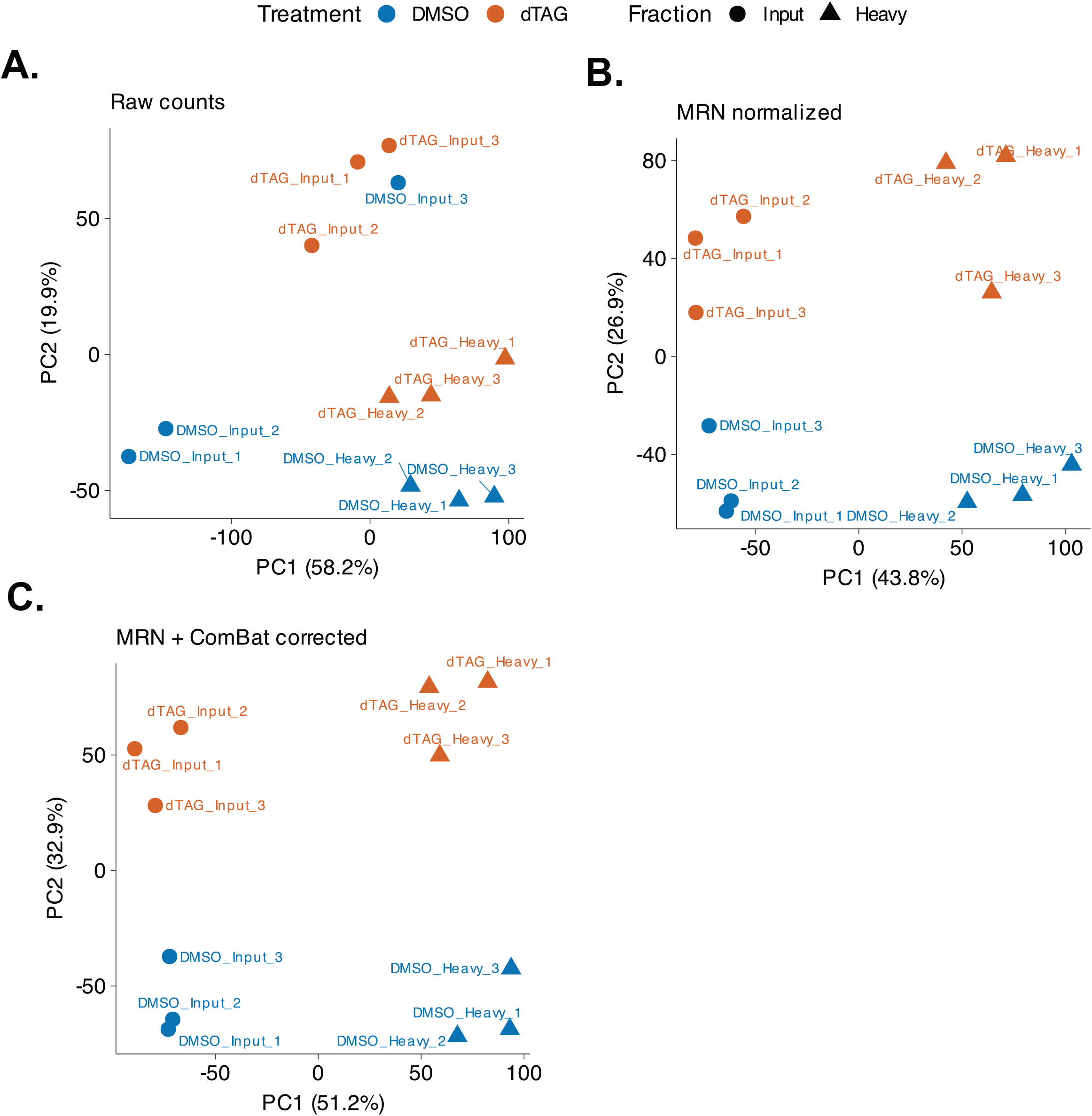
Principal component analysis validates ComBat batch correction. PCA of (A) raw gene-level counts, (B) DESeq2 median-ratio normalized counts, and (C) normalized, ComBat-corrected counts, colored by treatment (DMSO, dTAG) and shaped by fraction (Input, Heavy), for all three biological replicates. Batch-associated separation present in the raw and normalized-only data is resolved after ComBat correction, with samples clustering primarily by treatment and fraction rather than by replicate. Percentages indicate variance explained by each principal component.

**Figure. S2.**
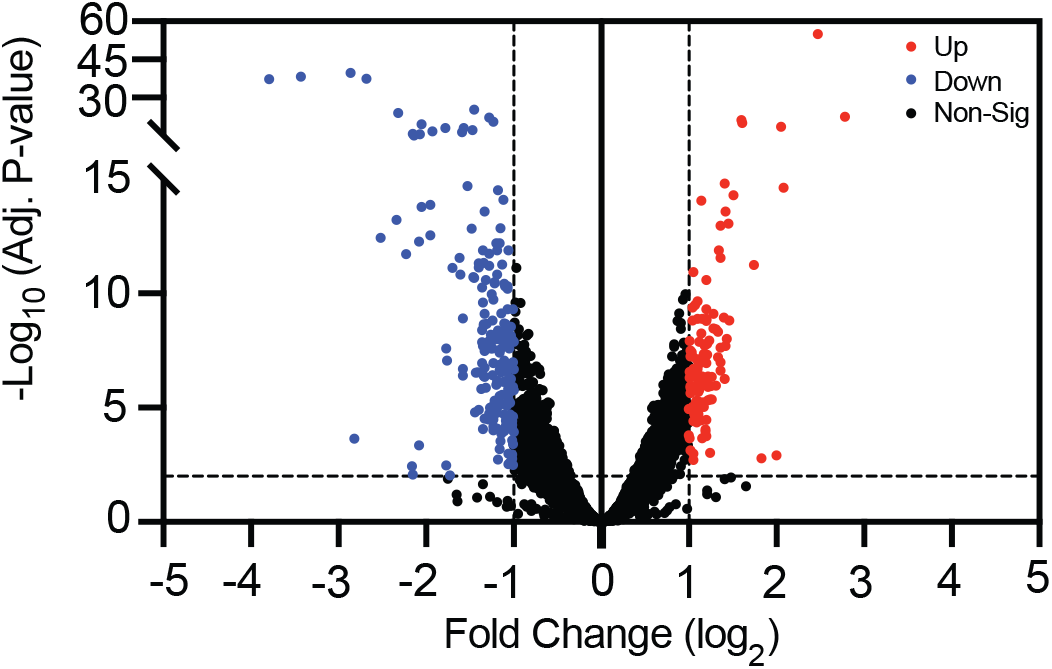
Minimal transcriptome remodeling upon acute EIF4G1 depletion. DESeq2 volcano plot of input RNA-seq (total mRNA) comparing dTAG-treated versus DMSO-treated cells. The minimal number of differentially expressed genes confirms that changes in polysome association (RPA) reflect translational redistribution rather than transcriptional remodeling.

**Figure. S3.**
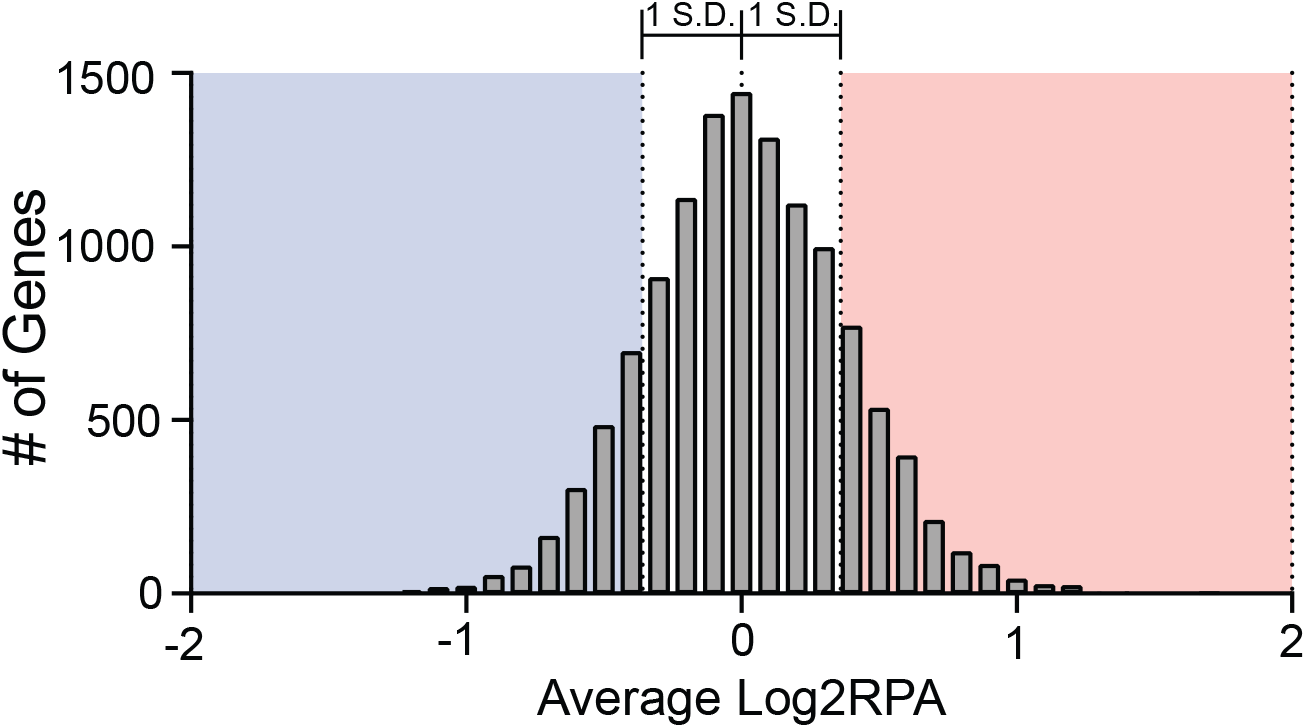
Distribution of log₂RPA values and classification threshold. Distribution of mean log₂RPA values across all quantified genes. Genes were classified as Resistant or Sensitive if their mean log₂RPA exceeded one standard deviation from the mean of the distribution in either direction (±0.38, corresponding to log₂(1.3), or approximately a 30% shift in RPA), and this shift was statistically significant by a moderated one-sample t-test (limma) with Benjamini-Hochberg FDR < 0.05. The ±0.38 cutoff is indicated.

**Figure. S4.**
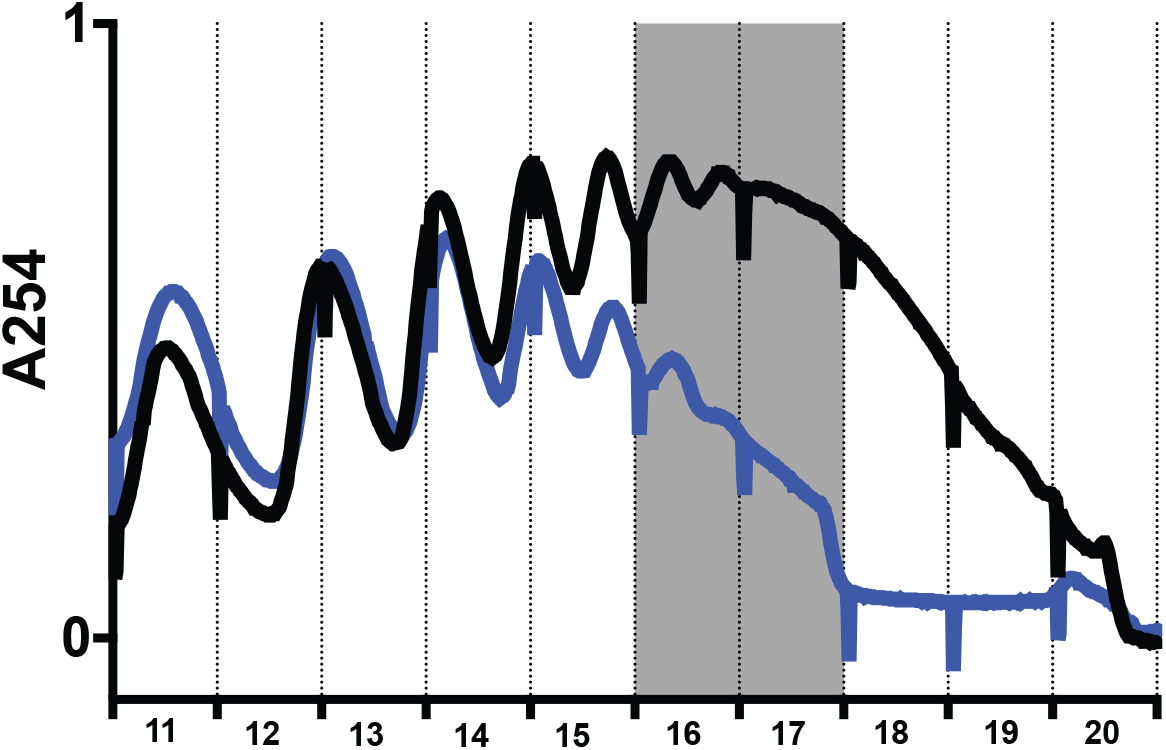
Polysome traces with fraction boundaries used for RNA-seq heavy polysome pooling. (A) Sucrose gradient polysome profiles showing the fraction boundaries used to define the heavy polysome pool (≥3 ribosomes per transcript) for RNA-seq library preparation. Fractions to the right of the indicated boundary were pooled as the heavy polysome fraction for biological replicates 1-3 (B-D).

**Figure. S5.**
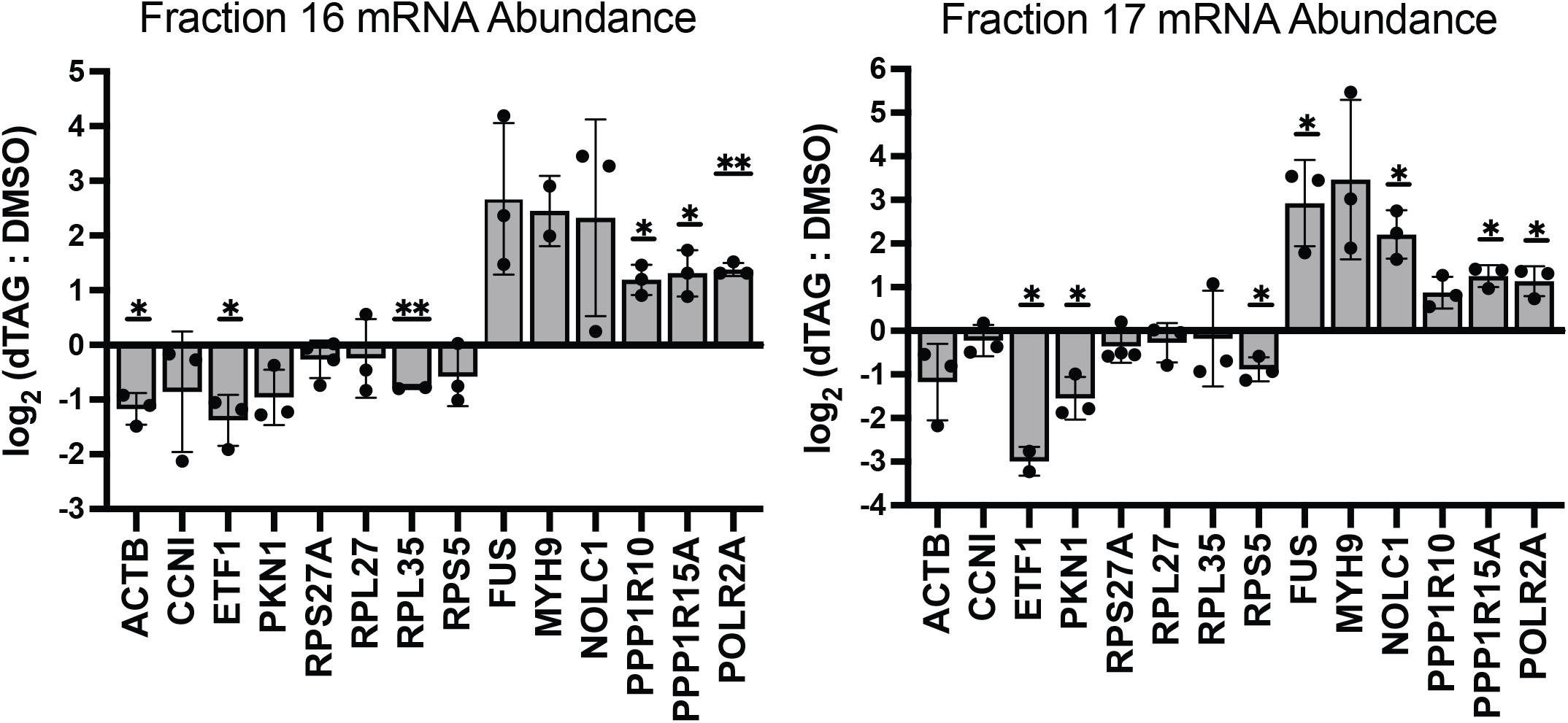
Per-fraction analysis of Resistant and Sensitive transcripts across heavy polysome window. Per-fraction qPCR log₂ fold changes (dTAG/DMSO) for representative Sensitive and Resistant transcripts across individual heavy polysome fractions (fractions 16–17).

**Figure. S6.**
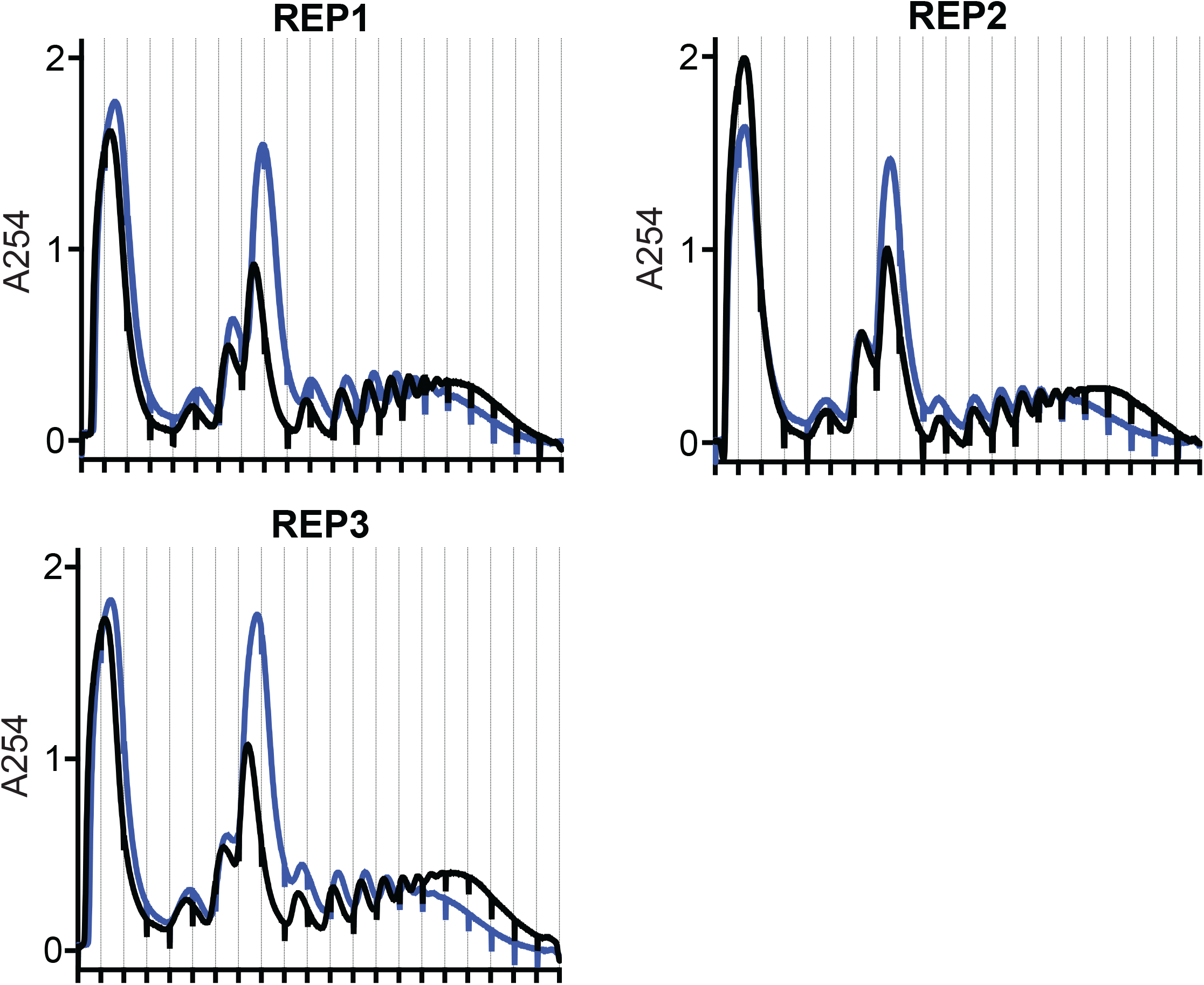
Polysome gradient profiles are consistent across biological replicates. Absorbance (A254) traces from sucrose gradient fractionation are shown for three independent biological replicates (REP1, REP2, REP3). Black traces represent DMSO-treated (vehicle control) cells and blue traces represent dTAG-treated cells.

**Figure. S7.**
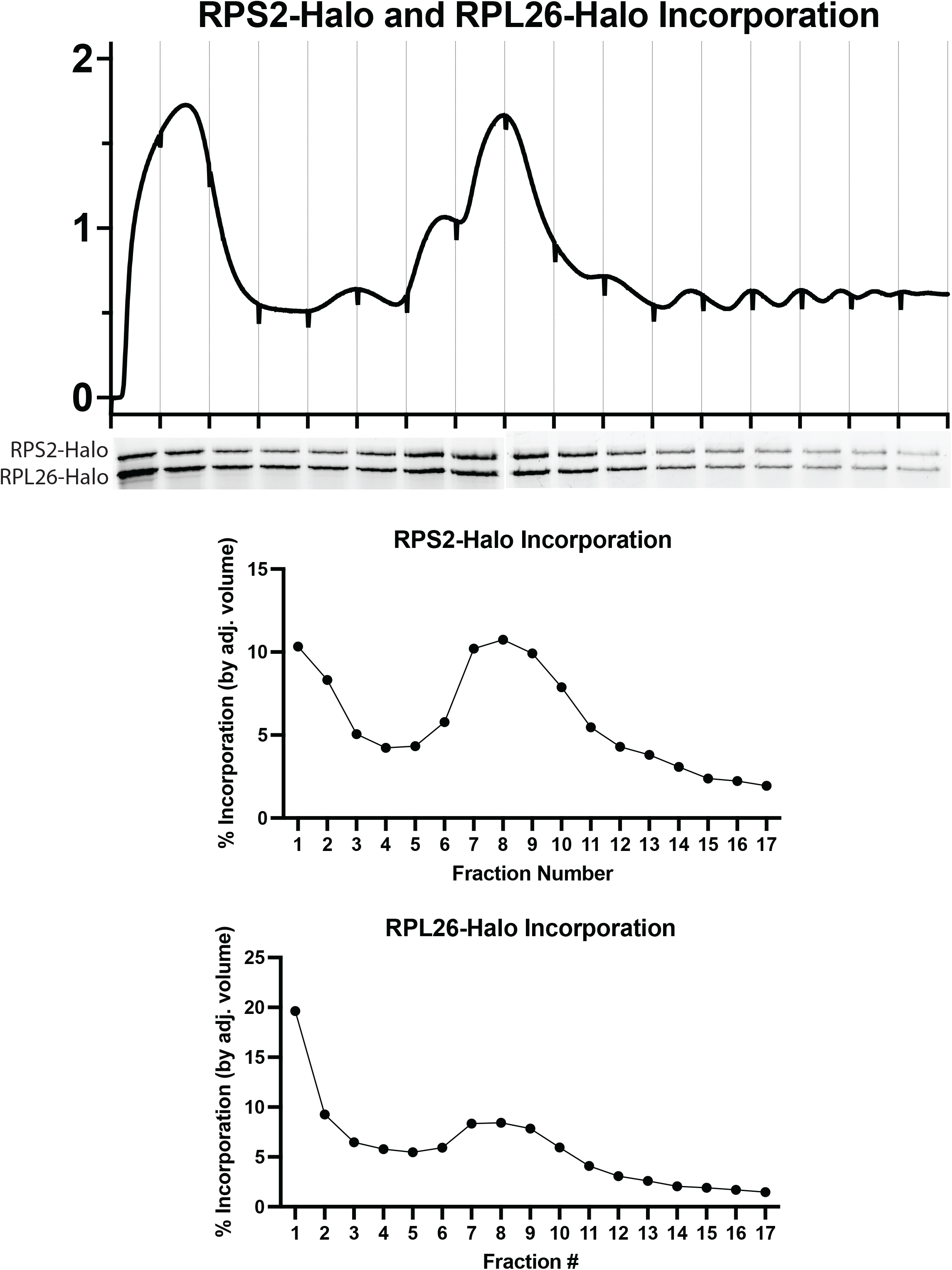
RPS2-Halo and RPL26-Halo predominantly incorporate into ribosomal subunits and fully assembled ribosomes. Sucrose gradient polysome trace with corresponding RPS2-Halo and RPL26-Halo in-gel fluorescence signal across the gradient, and quantification of RPS2-Halo and RPL26-Halo incorporation (percent of total signal) across fractions 1-17.

**Figure. S8.**
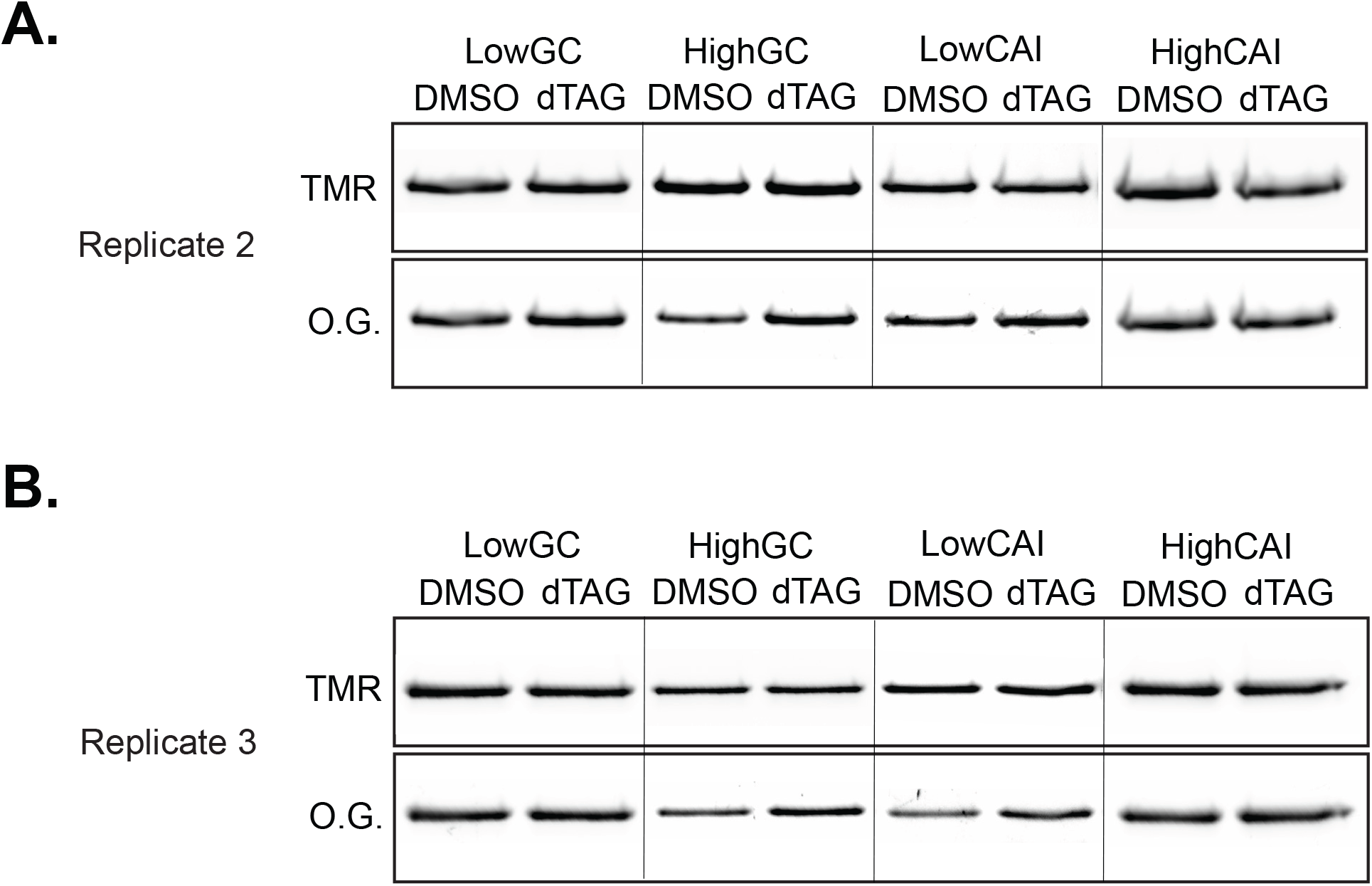
Biological replicates 2 and 3 of the CDS GC content and codon optimality Halo reporter experiment. (Low-GC, High-GC, Low-CAI, High-CAI constructs; DMSO vs. dTAG). SDS-PAGE fluorescent gel images showing TMR (preexisting protein) and Oregon Green (newly synthesized protein) signals for all four constructs under DMSO and dTAG conditions, for (A) biological replicate 2 and (B) biological replicate 3, corresponding to the quantification in Figure 5E.

**Figure. S9.**
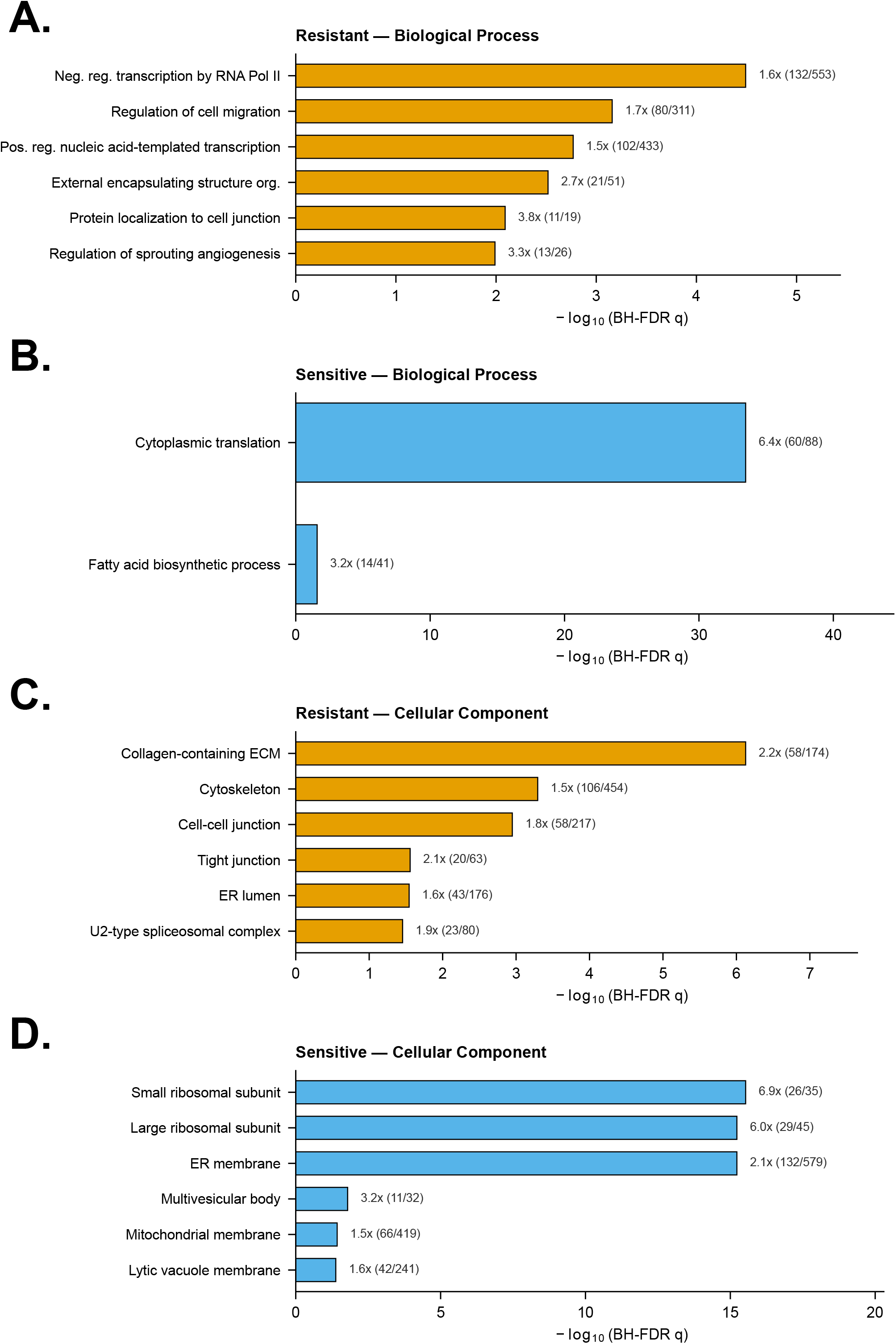
Gene Ontology enrichment among Resistant and Sensitive transcripts. (A) GO Biological Process enrichment among Resistant transcripts (hypergeometric test, Benjamini-Hochberg FDR-corrected; top terms by q-value shown). (B) GO Biological Process enrichment among Sensitive transcripts, same method. (C) GO Cellular Component enrichment among Resistant transcripts, same method. (D) GO Cellular Component enrichment among Sensitive transcripts, same method.

**Figure. S10.**
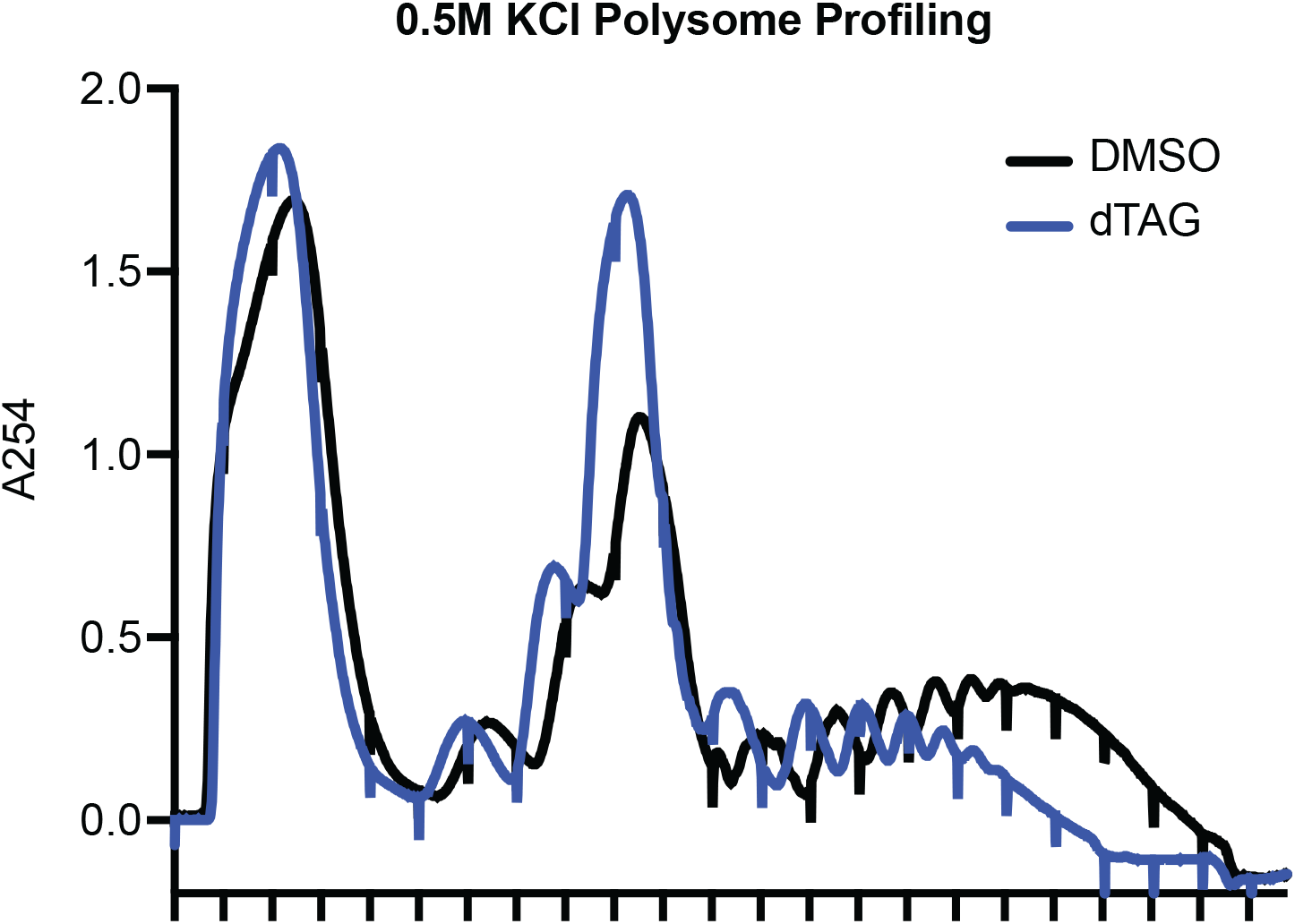
The 80S accumulation upon EIF4G1 depletion persists at high salt. Sucrose gradient polysome profiles (absorbance at 254 nm) of cells treated with DMSO or dTAGV-1 and fractionated in 500 mM KCl.

## Supplemental Tables (excel document)

Supplemental Table S1: Quantification of polysome metrics (monosome/polysome areas, P/M ratios).

Supplemental Table S2: Transcriptionally upregulated/downregulated genes (DESeq2).

Supplemental Table S3: EIF4G1 RPA Classification Summary Metrics

Supplemental Table S4: Antibodies used.

Supplemental Table S5: RNA-seq read filtering and mapping metrics.

Supplemental Table S6: qPCR primers.

Supplemental Table S7: 5’ Terminal Oligopyrimidine (TOP) mRNAs in EIF4G1 Dataset

Supplemental Table S8: Codon Adaptation Index (CAI) by classification group (Non-significant, Sensitive, Resistant, DAP5-Dependent).

Supplemental Table S9: HaloTag reporter constructs used in the GC content and codon optimality reporter assays (Figure 5).

Supplemental Table S10: Relative protein and mRNA abundance of the eIF4G paralogs EIF4G1, EIF4G2 (DAP5), and EIF4G3 in HCT116 cells.

